# CSPG4-targeting CAR-macrophages inhibit melanoma growth

**DOI:** 10.1101/2024.06.04.597413

**Authors:** Daniel Greiner, Qian Xue, Trinity QA Waddell, Elena Kurudza, Rachel L Belote, Gianpietro Dotti, Robert L Judson-Torres, Melissa Q Reeves, Samuel H Cheshier, Minna Roh-Johnson

**Affiliations:** Department of Biochemistry, University of Utah School of Medicine; Salt Lake City, UT, 84112, USA; Department of Neurosurgery, Clinical Neurosciences Center, University of Utah; Salt Lake City, UT, 84112, USA; Department of Dermatology, University of Utah School of Medicine; Salt Lake City, UT, 84112, USA; Department of Molecular Genetics, The Ohio State University College of Arts and Sciences; Columbus, OH, 43210, USA; Department of Microbiology and Immunology, University of North Carolina, Chapel Hill, NC 27599, USA; Department of Oncological Sciences, University of Utah School of Medicine; Salt Lake City, UT, 84112, USA; Huntsman Cancer Institute, University of Utah School of Medicine; Salt Lake City, UT, 84112, USA; Department of Pathology, University of Utah School of Medicine; Salt Lake City, UT, 84112, USA; Division of Pediatric Neurosurgery, Intermountain Primary Children’s Hospital; Salt Lake City, UT, 84112, USA

**Keywords:** Chimeric antigen receptor, macrophage, melanoma, anti-CD47, phagocytosis

## Abstract

Chimeric antigen receptor (CAR) T-cell therapy has revolutionized the treatment of hematological malignancies but has been clinically less effective in solid tumors. Engineering macrophages with CARs has emerged as a promising approach to overcome some of the challenges faced by CAR-T cells due to the macrophage’s ability to easily infiltrate tumors, phagocytose their targets, and reprogram the immune response. We engineered CAR-macrophages (CAR-Ms) to target chondroitin sulfate proteoglycan 4 (CSPG4), an antigen expressed in melanoma, and several other solid tumors. CSPG4-targeting CAR-Ms exhibited specific phagocytosis of CSPG4-expressing melanoma cells. Combining CSPG4-targeting CAR-Ms with CD47 blocking antibodies synergistically enhanced CAR-M-mediated phagocytosis and effectively inhibited melanoma spheroid growth in 3D. Furthermore, CSPG4-targeting CAR-Ms inhibited melanoma tumor growth in mouse models. These results suggest that CSPG4-targeting CAR-M immunotherapy is a promising solid tumor immunotherapy approach for treating melanoma.

**STATEMENT OF SIGNIFICANCE:** We engineered macrophages with CARs as an alternative approach for solid tumor treatment. CAR-macrophages (CAR-Ms) targeting CSPG4, an antigen expressed in melanoma and other solid tumors, phagocytosed melanoma cells and inhibited melanoma growth *in vivo*. Thus, CSPG4-targeting CAR-Ms may be a promising strategy to treat patients with CSPG4-expressing tumors.

## INTRODUCTION

Chimeric antigen receptor (CAR) T-cell therapy has demonstrated the groundbreaking potential to harness the immune system against cancer. While this revolutionary approach has reshaped the treatment of hematological malignancies, its promise has yet to be fully realized in the context of solid tumors (1). Several complexities specific to solid tumor biology present unique obstacles that have limited the success of CAR-T cell therapies. Tumor heterogeneity presents a formidable challenge for CAR-T cell therapy. Antigen heterogeneity in solid tumors means that CAR-T cells designed to target a single antigen may only eliminate cells with high antigen expression, leaving behind populations with low or no expression, resulting in therapeutic resistance and tumor relapse (2). Solid tumors also have dense physical barriers of extracellular matrix that impede active T-cell infiltration (3) and cultivate a highly immunosuppressive microenvironment. This environment is characterized by the presence of suppressive immune cells (like regulatory T cells, myeloid-derived suppressor cells, and tumor-associated macrophages) (4,5), inhibitory cytokines (such as TGF-beta) (6), hypoxia (7,8), and metabolic factors that can disarm and exhaust CAR-T cells, ultimately limiting their anti-tumor activity (9). While efforts to further engineer CAR-T cells for improved activity against solid tumors are ongoing, an alternative approach is engineering other immune cells, like macrophages that inherently overcome these challenges.

Unlike T-cells, macrophages comprise a large portion of the tumor mass in solid tumors (5). Macrophages are professional phagocytes, eating their targets and easily infiltrating barriers surrounding solid tumors (10). Macrophages can also reprogram other immune cells such as regulating the polarization of other macrophages (11) and educating adaptive immune cells through antigen presentation (10). Given that macrophages phagocytose their tumor targets, the antigen presentation capabilities of macrophages suggest that macrophages will present multiple tumor antigens to T-cells, thus potentially bypassing challenges associated with tumor heterogeneity. The central role that macrophages play in the immune system suggests that macrophages are an ideal cell to use for adoptive cell therapies in cancer, as reprogramming the tumor immune response is likely to lead to a sustained and robust anti-tumor response.

Several groups have explored the potential of using chimeric antigen receptor-expressing macrophages (CAR-Ms) to treat various types of solid tumors such as ovarian cancer. These CAR-Ms have predominantly targeted a variety of canonical tumor antigens including, but not limited to, CD19, CD47, HER2, and EGFRvIII (11–25). These studies show that CAR-Ms targeting certain cancer types are specific and efficient, and CAR-Ms effectively reduce tumor burden in *in vivo* xenograft models. However, it is still unclear whether CAR-Ms will be an effective strategy to treat all solid tumors, and whether the ideal cancer antigen has been selected. Thus, we sought to engineer CAR-Ms against a melanoma-specific antigen to treat metastatic melanoma.

Cutaneous melanoma is a form of skin cancer that arises from melanocytes. According to the latest Surveillance, Epidemiology, and End Results (SEER) data, for patients who are diagnosed with distant metastatic melanoma, the 5-year survival is below 40% (26). In the last 10 years, immune checkpoint blockade (ICB) approaches have drastically improved the survival of patients with metastatic melanoma. However, despite this remarkable success, approximately 50% of patients with metastatic melanoma do not benefit from ICB, even with combinations of αCTLA-4 and αPD-1 (27). Patients can exhibit both primary resistance and acquire resistance during treatment (27). Thus, it remains essential to explore additional treatment options, particularly for patients who do not respond to current treatment strategies.

We sought to develop CAR-Ms that recognize and phagocytose melanoma cells. Numerous tumor-associated antigens enriched in melanomas have been identified, and we focused on chondroitin sulfate proteoglycan 4 (CSPG4), also known as NG2 and HMW-MAA (28). CSPG4 is a ∼300 kDa transmembrane proteoglycan that regulates cancer cell migration, invasion, epithelial-mesenchymal transition, and proliferation (29). CSPG4 is frequently highly expressed in melanoma tumors, and expression in non-malignant cells is low, making it an ideal target antigen (29,30). Furthermore, a CAR-T cell therapy targeting CSPG4 is currently being used in a phase I clinical trial for head and neck cancer (NCT06096038) (31), suggesting CSPG4-targeting CARs are specific with minimal off-target effects.

In this study, we demonstrate that CSPG4-targeting CAR-Ms phagocytose melanoma cells *in vitro,* and CSPG4-targeting CAR-Ms in combination with CD47 blocking antibodies efficiently inhibit melanoma spheroid growth in 3D. Furthermore, we show that CSPG4-targeting CAR-Ms inhibit melanoma growth *in vivo*. Thus, this work provides a potential therapy for melanoma patients.

## MATERIALS AND METHODS

Detailed information on all key resources is available in Supplemental Table 1.

### Cell culture

A375, YUMM1.7, B16F10 cell lines were obtained from the ATCC. Short-tandem repeat confirmed 624-mel and WM793 cell lines were obtained from the Judson-Torres lab at the University of Utah. YUMM1.1, YUMM3.2, and YUMM5.2 cells were generously provided by Matthew Williams’ lab at the University of Utah. A375 cells were cultured in DMEM (Thermofisher #11965118) supplemented with 10% FBS. 624-mel and WM793 cells were cultured in RPMI (Thermofisher #11875119) supplemented with 10% fetal bovine serum (FBS) (Sigma Aldrich #F4135) and 1% Glutamax (Thermofisher #35050061). YUMM1.1, YUMM1.7, YUMM3.2, and YUMM5.2 cells were cultured in DMEM/F12 (Thermofisher #11330057) supplemented with 10% FBS and 1% MEM non-essential amino acids (Thermofisher #11140050). Cell lines were routinely tested for mycoplasma detection using a PCR detection kit (ATCC, #30-1012K). Cells were used for <20 passages.

### Lentiviral particle production

pCMV-VSV-G, psPAX2, and transgene plasmids were transfected into HEK293FT cells to generate lentivirus media as previously described (33) and concentrated using Lenti-X concentrator (Takara Bio #631232) according to the manufacturers protocol.

### Generation of fluorescent tumor cell lines

Lentivirus containing pLKO-Lck-mScarlet or pLenti6-H2B-mCherry was generated from HEK293FT cells as described in ‘Lentiviral particle production’. To generate shCSPG4 cell lines, lentivirus containing plasmids from either non-target short hairpin RNA (Sigma #SHC0020) or shCSPG4 (gene target NM_001897) (Sigma #TRCN0000422139 or Sigma #TRCN0000437747), psPAX2, and pCMV-VSV-G were generated from HEK293FT cells as described in ‘Lentiviral particle production’. All short hairpin RNA constructs were cloned into the pLKO.1 backbone. Melanoma cells expressing Lck-mScarlet were transduced using concentrated lentivirus containing the short hairpin RNA.

### Human primary cell isolation and CAR generation

Anonymous donor healthy human blood in either leukoreduction filters or apheresis cones were obtained from Associated Regional and University Pathologists, Inc (ARUP). Leukocytes were recovered from blood, CD14+ monocyte isolated by adhesion, transduced, and cultured as previously described (33).

### Mouse primary cell isolation and CAR generation

Bone marrow monocyte-derived macrophages (BMDMs) were isolated from C57BL/6 mice femurs as previously described (34). BMDMs were cultured in RPMI supplemented with 10% FBS, 1% Penicillin-Streptomycin, 1% Glutamax, 1% Sodium Pyruvate (Thermofisher #11360070), 0.1% 2-Mercaptoethanol and 50 ng/mL recombinant human M-CSF (Peprotech #300-25). BMDMs were transduced with concentrated lentivirus on day 3 with subsequent media changes every 2 days.

### 2D Phagocytosis flow assay

Cells were detached, counted, and CAR-Ms were plated with melanoma cells for 24 hours. Cells were detached and processed for flow cytometry. CAR-M phagocytosis percentages were determined by gating live Cells, singlets gates, CD11B+ macrophages, GFP+ for transduced macrophages, and setting an Lck-mScarlet+ / GFP+ gate for phagocytosis based on 0-hour coculture (cells pooled together just prior to flow). See Supplemental Fig. 2D for full gating strategy.

### 2D Phagocytosis imaging assay

A375-Lck-mScarlet cells and CAR-Ms were plated together as described in 2D Phagocytosis flow assay and cultured for 24 hours. Images were acquired on a Zeiss LSM 880 Airy Scan microscope using either the LSM acquisition mode or Airyscan FAST acquisition mode (and subject to deconvolution using Zen software (Carl Zeiss)) with ‘auto’ settings. Maximum intensity projections of images or a central z-plane were analyzed in FIJI (35) (version 2.14.0) such that an automated mask was generated around GFP+ macrophages. Lck-mScarlet signal intensity was measured inside the GFP+ mask. Lck-mScarlet signal surface area was manually masked on maximum intensity projection images (Supplemental Fig. 3E). For images showing x-z, or y-z representations (Supplemental Fig. 3A, 6A), images were resliced in FIJI, and then scaled on the z-axis to improve visibility.

### 2D Timelapse overnight imaging

A375-Lck-mScarlet cells and CAR-Ms were plated together as described in ‘2D Phagocytosis imaging assay’ and cultured for 1 hour to allow the cells to settle. Images were acquired using Airyscan FAST acquisition mode every 10 minutes for 18 hours and maintained at 37°C and at 5% CO_2_ with an on-stage incubator.

### 3D Phagocytosis flow assay

CAR-Ms were plated with Lck-mScarlet or H2B-mCherry transduced melanoma cells for 24 or 72 hours to generate spheroids with 10 µg/mL of IgG control antibody or αCD47. Four spheroids were pooled for each technical replicate, then pelleted, and the supernatant removed. Cells were processed for flow cytometry as previously described in ‘2D Phagocytosis flow assay’.

### Nuclei status assay

Spheroids were prepared as described in ‘3D Phagocytosis flow assay’. Spheroids were then dissociated into single cells by pipetting and plated on glass bottom imaging dishes for 6 hours for cells to adhere. Cells were fixed, then permeabilized and stained with DAPI. Images were acquired on a Zeiss LSM 880 microscope using LSM acquisition mode. H2B-mCherry cells were manually counted for either unengulfed (A375-H2B-mCherry cell not surrounded by GFP+ macrophages), engulfed and live (A375-H2B-mCherry surrounded by GFP+ macrophage in X, Y, and Z plane, with H2B-mCherry colocalizing with DAPI), or engulfed and dead (A375-H2B-mCherry surrounded by GFP+ macrophage in X, Y, and Z plane, with H2B-mCherry signal dispersed and not colocalizing with DAPI).

### 3D Spheroid growth assay

A375-H2B-mCherry cells or YUMM1.7-H2B-mCherry cells were plated for 72 hours to generate spheroids. CAR-Ms or control CAR-Ms were then added to the wells along with 10 µg/mL of IgG control antibody or αCD47 for 10 days. Media was changed every 3 days by removing 100 µL of media and replacing it with fresh media and treatments. Spheroids were imaged every 8 hours using an Incucyte SX5 analysis system, and H2B-mCherry total object integrated intensity and largest object area were quantified using the spheroid analysis module following spectral unmixing using Incucyte software (version 2020C Rev1).

### CAR-M adherence assay

A375-Lck-mScarlet cells were plated as described in ‘3D spheroid growth assay’. CAR-M were added to the wells along with 10 µg/mL of IgG control antibody or αCD47 for 8 hours. The spheroids were then physically removed from the well with a wide-bore pipette tip, mixed with Matrigel, and then added to imaging dishes. Images were then acquired of the spheroids with on a Zeiss LSM 880 microscope using Airyscan FAST acquisition mode. Images were subject to deconvolution using Zen software with ‘auto’ settings and then processed in FIJI to generate a mask of the Lck-mScarlet+ spheroid. The number of GFP+ cells that were attached to the spheroid was then quantified.

### CSPG4 expression flow assay

Cells were plated and incubated for 72 hours. Cells were then scraped into fresh media and then pelleted at 300G for 5 minutes. Cells were stained with αCSPG4 antibodies or isotype control antibodies and secondary antibodies on ice, protected from light. Cells were then washed with PBS, then resuspended in an appropriate volume of flow buffer and analyzed for fluorescence expression on the Fortessa. Cells were analyzed for live cells, singlets as described in ‘2D Phagocytosis Flow Assay’ and then gated for CSPG4+ cells against isotype-stained control cells.

### Image cytometry

Cells were plated as described above in 2D Phagocytosis flow assay (Fig. 3B, C) or 3D phagocytosis flow assay (Supplemental Fig. 5A). After 24 hours of coculture, cells were prepared for flow as previously described in ‘2D Phagocytosis flow assay’. Cells were stained with DAPI for viability gating. Imaging cytometry was performed on an Imagestream Mk II (Amnis). Cells were gated by Area/Aspect ratio, in-focus, DAPI for viability, total GFP+ events, non-saturated GFP+ events. Internalization events were detected by setting a minimum RFP+ intensity and applying a GFP signal adaptive erode to detect an RFP+ signal within a GFP+ signal. Internalized phagocytosis events were classified by size based on the area intensity of the RFP+ signal (∼75-microns).

### Xenograft experiments

Animal experiments were approved by the Institutional Animal Care and Use Committee (IACUC) at the University of Utah and conducted with assistance from the Preclinical Research Shared Resource at Huntsman Cancer Institute Research. 250K A375-H2B-mCherry cells were injected with Matrigel subcutaneously into 6-8 week-old female NOD.Cg-*Rag1^tm1Mom^Il2rg^tm1Wjl^*/SzJ (NRG) mice. The subsequent injections were performed peri-tumorally with vehicle and CAR-Ms only in PBS on days indicated. For same-day injections only, at first injection, vehicle or CAR-Ms were injected with the tumor cells in Matrigel. Tumor volume measurements were taken with calipers as indicated on graphs until experimental endpoints.

### Tumor digestion for flow cytometry

Tumors were mechanically dissociated with scalpels, then digested with RPMI (+.032% Aqueous Collagenase D, +.008% DNAse) for 40 minutes at 37°C on a shaker, inverting every 10 minutes. Digested tumor was then strained over 70-micron filters, washed in PBS, and then cells were counted for flow cytometry.

### Tumor immunostaining and imaging

Tumors were harvested and fixed in formalin prior to embedding in paraffin. Samples were deparaffinized and rehydrated using Citrisolv (VWR #89426-268), followed by rinses in sequential dilutions (100%, 95%, 80%, 70%) of ethanol, and then ddH_2_O and PBS. Antigen retrieval was done in Tris-EDTA (pH 9) buffer overnight at 60°C, and then washed 2x with ddH_2_O and PBS. Samples were blocked 1 hour, followed by primary chicken anti-GFP (ABCAM #13970) (1:500) for 1 hour, all at room temperature. Samples were washed in TBS-T, and then stained for 1 hour with Goat Alexa fluor 488 anti-Chicken (1:500) (Jackson Immuno #103-545-155) at room temperature. Autofluorescence was then quenched using 0.1% Sudan Black in 70% ethanol for 10 minutes at room temperature, and then stained with DAPI prior to mounting and imaging. Images were acquired with a Zeiss LSM880 microscope using LSM acquisition mode. We used E0771 tumors grown in mice expressing GFP-tagged mitochondria in macrophages (mice carrying both of the following transgenes: B6.Cg-Gt(ROSA)26Sortm1(CAG-EGFP)Brsy/J (Jackson Laboratory #032290) and a B6.129P2-Lyz2tm1(cre)Ifo/J (Jackson Laboratory #004781)) to set a threshold for positive GFP expression.

### Image analysis

All images were acquired with a Zeiss LSM 880 microscope or an Incucyte SX5 system. For images acquired with the LSM 880, selected z-planes were used to generate maximum intensity projections using Zen software. Linear adjustments to brightness and contrast were done with FIJI. Representative images were cropped and assembled with Adobe Photoshop (version 24.4.1) and Illustrator (version 27.5).

### Bioinformatic analysis

CSPG4 normalized expression values from fresh resected healthy human skin and metastatic melanoma specimens were obtained from previously published single-cell RNA-sequencing (scRNAseq) data sets GSE151091 and the Single Cell portal (https://portals.broadinstitute.org/single_cell/study/melanoma-immunotherapy-resistance) (36,37). Both studies were analyzed using smartseq 2 based pipelines and then compared using rank mean normalization. Only non-cycling healthy human skin cells were included in the analysis. Immunotherapy response data was obtained as in Riaz et al (38), via the Immuno-genomic atlas for immune checkpoint blockade-based cancer immunotherapy(http://bioinfo.vanderbilt.edu/database/Cancer-Immu/) (39). Data visualization and statistical analyses were performed in GraphPad Prism (version 9.4.1), python (version 3.7.4), and R (version 4.3.2).

### Statistical analysis

All statistical analysis was done using GraphPad Prism and presented as mean values +/- standard error of the mean (SEM). Outliers greater than two standard deviations were removed from analysis. The figure legend indicates the statistical test used and the number of biological and technical replicates. To determine the number of biological replicates, we sought to detect a 50% difference in phagocytosis between conditions, assuming a 10% standard deviation. Thus, we required a minimum of 3 biological replicates for our studies. Flow cytometry data were analyzed using FlowJo software (version 10.10.0). Because animal experiments were not performed across both male and female mice, we are not addressing sex as a biological variable in these studies.

## RESULTS

### CSPG4-targeting CAR-Ms phagocytose CSPG4-expressing cancer cells

CSPG4 was first characterized as overexpressed in melanoma; however, CSPG4 upregulation in numerous other cancers including breast and glioblastoma has since been described (28,40). CSPG4 has been reported to promote tumor growth and survival through numerous signaling pathways associated with its extracellular domains (40). Transcriptional analysis suggests that CSPG4 mRNA expression is low in healthy tissue, with undetectable protein expression levels by immunohistochemistry (30). We analyzed published single-cell RNA-sequencing data from fresh healthy human skin and metastatic melanomas (36,37,41). These analyses revealed significantly higher CSPG4 transcript abundance in a majority of malignant tumor cells from patient samples compared to healthy skin cells, including melanocytes (Fig. 1A; Supplemental Fig. 1A). Importantly, non-malignant cells, both in the tumor microenvironment (Supplemental Fig. 1B) and across broader cell types (Supplemental Fig. 1C), generally presented low CSPG4 transcript counts, with notable exceptions being Sertoli cells, oligodendrocyte precursors cells, and a population of smooth muscle cells. However, immunohistochemistry approaches do not reveal high levels of CSPG4 protein expression in oligodendrocyte precursor cells in adult brain (42), and the blood-testis barrier represent one of the tightest blood-tissue barriers in the body, protecting Sertoli cells (43). In pre-clinical studies of rats treated with a cross-reactive rat IgE antibody against CSPG4, rats did not exhibit gross abnormalities or long-term toxicity, suggesting that CSPG4-targeting strategies do not yield significant off-target effects (44).

**Figure 1.**
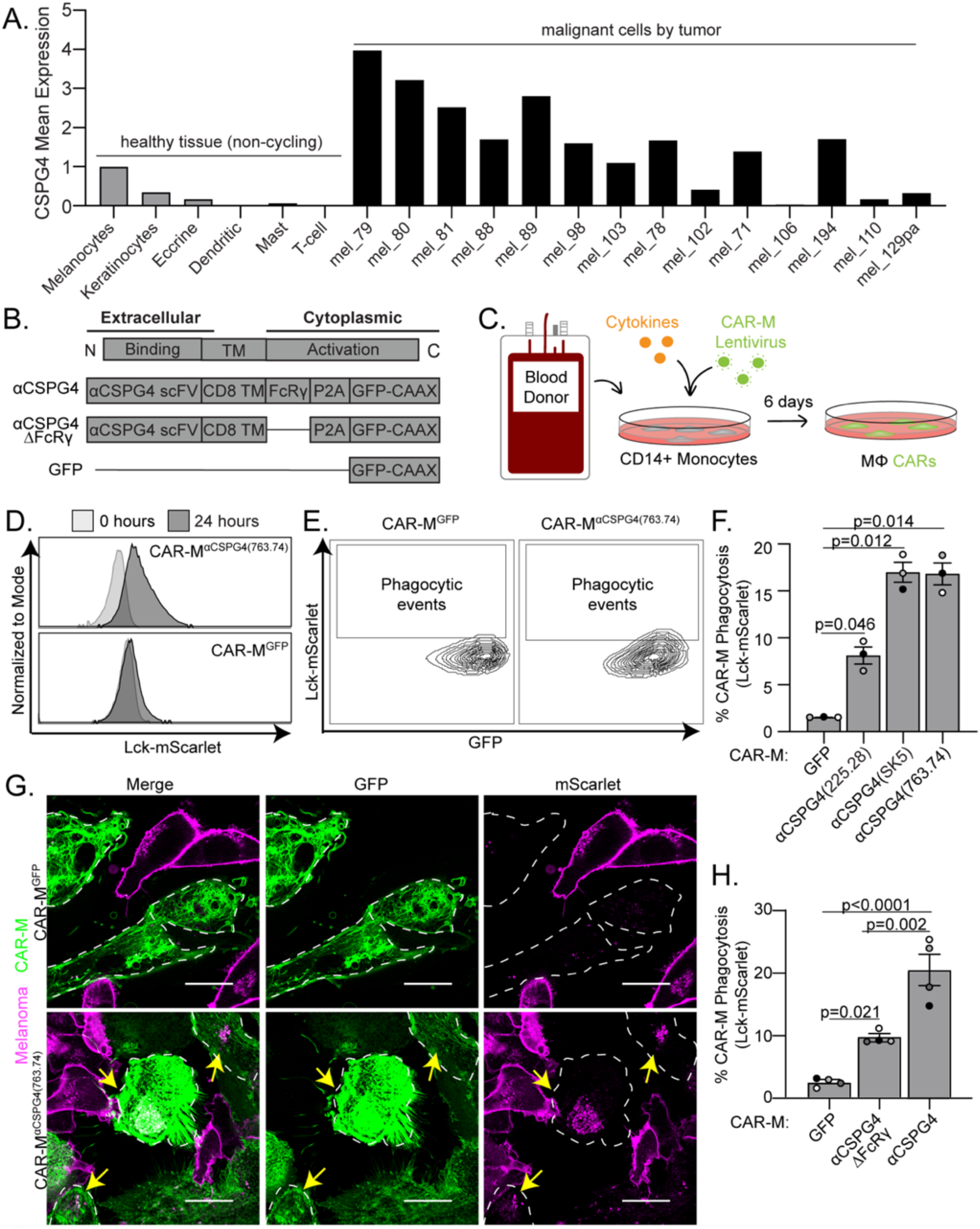
CSPG4-targeting CAR-Ms phagocytose CSPG4-expressing melanoma cells in 2D cell culture. **A)** CSPG4 transcript levels by single-cell RNA sequencing analysis of healthy (non-cycling) tissue vs melanoma tumors, rank mean normalized. **B)** Schematic of chimeric antigen receptor macrophage constructs. **C)** Schematic for generation of primary human CAR-Ms. **D)** Representative histograms showing phagocytic events after 24-hour coculture for CAR-M^GFP^ and CSPG4^αCSPG4(763.74)^. **E)** Representative flow cytometry contour plots showing CAR-M-mediated phagocytosis for CAR-M^GFP^ control and CSPG4^αCSPG4(763.74)^. **F)** Quantification of CAR-M-mediated phagocytosis shown in (D, E) for CAR-M^GFP^ and CAR-M with 3 scFvs (225.28, SK5, 763.74). **G)** Representative single z-plane images of CAR-M ^αCSPG4(763.74)^ (green)-mediated phagocytosis of A375-Lck-mScarlet cells (magenta) after 24 hours of 2D coculture by microscopy. White dashed line outlines CAR-M^αCSPG4^ cell boundary, yellow arrows indicate CAR-M phagocytosis. Scale bar is 20 microns. **H)** Quantification of CSPG4^αCSPG4(763.74)^ phagocytosis compared to CAR-M^GFP^ and CAR-M^αCSPG4(763.74)ΔFcRγ^ by flow cytometry. For all graphs, each dot on the graph is a biological replicate, each as a shade of gray. F, H) Mean +/- SEM, 1-way ANOVA with Tukey’s multiple comparisons test.

We next determined the correlation between ICB therapy outcomes and CSPG4 expression. Since ICBs targeting the PD-1/PDL-1 axis is the most common immunotherapy in melanoma patients, we analyzed the expression in αPD-1 non-responders. We found that patients who did not respond to αPD-1 treatment correlated with higher CSPG4 expression (Supplemental Fig. 1D). Furthermore, αPD-1 treatment itself did not change CSPG4 expression (Supplemental Fig. 1E). These results suggest that CSPG4 is a good target for melanoma patients who did not respond well to ICB treatment.

To target CSPG4-expressing cells, we engineered CAR constructs incorporating one of three distinct single-chain variable fragments (scFv) exhibiting affinity for CSPG4: 225.28, SK5, and 763.74, each of which recognizes different extracellular domains (30,32,45,46) (Supplemental Fig. 2A). We followed previous CAR designs (12) using an FcRγ signaling domain, and combined this domain with CSPG4 scFvs. As controls, we designed CARs lacking the intracellular FcRγ phagocytic signaling domain, as well as a CAR expressing GFP alone, to control for activation associated with transduction (Fig. 1B). To generate CAR-Ms, we used human primary blood monocyte-derived macrophages, which we isolated from healthy blood donors, transduced monocytes with CSPG4-targeting CARs, and then differentiated the monocytes into macrophages (Fig. 1C). Using our previously established protocols (33), CSPG4-CAR expressing macrophages were generated with high transduction efficiencies (Supplemental Fig. 2B), and thus we proceeded with using primary macrophage-derived CAR-Ms for all subsequent studies.

We used the human metastatic melanoma cell line A375 to test whether CSPG4-targeting CAR-Ms phagocytose melanoma cells. A375 cells exhibit high CSPG4 expression, compared with primary macrophages from multiple donors as a negative control (Supplemental Fig. 2C). We engineered A375 cells to express a fluorescent membrane marker (Lck-mScarlet) to visualize the melanoma cells. We then cocultured A375-Lck-mScarlet cells with each of the three different CSPG4-targeting CAR-Ms. Using flow cytometry to quantify CAR-M-mediated phagocytosis as the percentage of CAR-Ms with A375-Lck-mScarlet fragments (Supplemental Fig. 2D), we found that all CAR-M^αCSPG4(225.28/SK5/763.74)^ phagocytosed melanoma cells at higher rates than the control CAR-M^GFP^ (Fig. 1D-F). These differences in phagocytosis rates were not due to the level of CAR transduction efficiency, as the percent CAR transduction efficiency in macrophages did not affect the percent of CAR-M-mediated phagocytosis (Supplemental Fig. 2E). To confirm these flow cytometric events were *bona fide* CAR-M-mediated phagocytosis, rather than A375 membrane fragments adhering to the exterior of CAR-Ms, we imaged CAR-M and A375 cocultures with high-resolution microscopy. After 24 hours of coculture, we observed Lck-mScarlet punctae inside of CAR-M^αCSPG4(225.28/SK5/763.74)^ cells compared to CAR-M^GFP^ (Fig. 1G; Supplemental Fig. 2F). We proceeded to use CAR-M^αCSPG4(763.74)^ (herein referred to as CAR-M^αCSPG4^) for the remainder of our studies due to the consistency of phagocytosis, availability of published data regarding its binding region on CSPG4 (47), and its current use in phase I CAR-T cell clinical trials (NCT06096038) (31). To better understand CAR-M-mediated phagocytosis of melanoma cells, we used timelapse recording and 3D visualization approaches. Using 3D reconstruction, we confirmed that the Lck-mScarlet punctae were fully internalized in the CAR-M^αCSPG4^ (Supplemental Fig. 3A; Supplemental Video 1). Live timelapse recordings reveal that CAR-M^αCSPG4^ began phagocytosing melanoma cells within the first 4 hours of coculture, with most events being trogocytosis (cell nibbling), rather than whole cell phagocytosis (Supplemental Fig. 3B; Supplemental Video 2).

We next tested whether the intracellular FcRγ region was critical for CAR-M-mediated phagocytosis. Consistent with previous studies with CD19-targeting CAR-Ms (12), removal of the FcRγ domain (αCSGP4ΔFcRγ) resulted in a statistically significant reduction in the percent phagocytosis by flow cytometry compared to the full CSPG4-targeting CAR-M (Fig. 1H). Interestingly however, CAR-M^αCSGP4ΔFcRγ^ exhibited higher phagocytosis rates than control CAR-M^GFP^ (Fig. 1H). We hypothesize that the physical interaction between the CSPG4-scFv and CSPG4 on the surface of melanoma cells is sufficient to promote CAR-M-mediated phagocytosis, likely via the maintenance of endogenous FcRγ signaling. Live cell imaging of CAR-M^αCSGP4ΔFcRγ^ cocultures with A375-Lck-mScarlet cells also show higher rates of phagocytosis of melanoma cells compared to CAR-M^GFP^ (Supplemental Fig. 3C, D). Taken together, these results suggest that CSPG4-targeting CAR-Ms efficiently phagocytose metastatic melanoma cells.

We next sought to determine the specificity of CSPG4-CAR-M-mediated phagocytosis. Using human melanoma cell lines with differing CSPG4 surface expression, we quantified the level of cancer cell phagocytosis by CAR-M^αCSGP4^ and CAR-M^GFP^. We found that A375 cells and WM793 melanoma cells exhibited high levels of CSPG4 expression on the surface of the cells, and 624-mel melanoma exhibited low CSPG4 expression (Fig. 2A, B). When we cocultured each cell line with control CAR-M^GFP^, we observed no differences in CAR-M-mediated phagocytosis. However, when we cocultured each of these cell lines with CAR-M^αCSPG4^, we observed high rates of phagocytosis with A375 and WM793 cells, but not with 624-mel cells (Fig. 2C). To further confirm specificity, we knocked down CSPG4 expression in A375 cells. We knocked down A375 surface expression with two different CSPG4 shRNAs to ∼40% compared to non-targeting controls (Fig. 2D, E) and found that the percent of CAR-M^αCSPG4^ phagocytosis was significantly reduced when CSPG4 expression was decreased on A375 cells compared to non-targeting controls (Fig. 2F). These results suggest that CAR-M^αCSPG4^ phagocytosis is specific to CSPG4-expressing cells.

**Figure 2.**
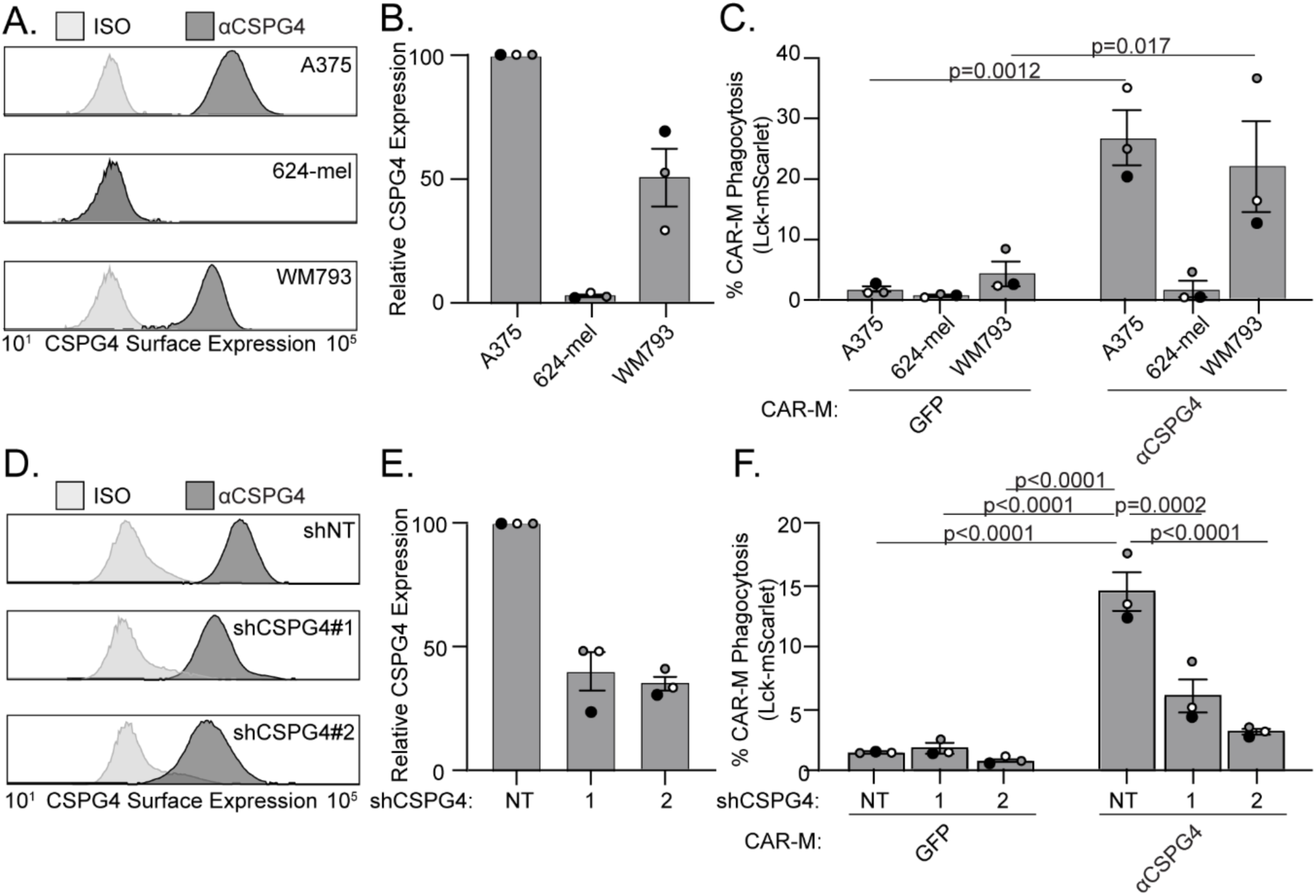
CSPG4-targeting CAR-M phagocytosis is specific to CSPG4-expressing cells. **A)** Representative flow plot of CSPG4 surface expression on A375, 624-mel, and WM793 cells with flow cytometry. **(B)** Quantification of CSPG4 surface expression in (A) normalized to A375 expression. **C)** Quantification of CAR-M-mediated phagocytosis (CAR-M^GFP^ and CAR-M^αCSPG4^) of A375, 624-mel, and WM793 cells after 24 hours of coculture. **D)** A representative flow plot of CSPG4 surface expression on A375 cells expressing non-targeted shRNA or 2 different shCSPG4 RNAs. **E)** Quantification of CSPG4 surface expression in (D) normalized to shNT expression. **F)** Quantification of CAR-M-mediated phagocytosis (CAR-M^GFP^ and CAR-M^αCSPG4^) of A375 cells treated with non-targeting shRNAs or 2 different shCSPG4 RNAs. For all graphs, each dot on the graph is a biological replicate, each as a shade of gray. C) Mean +/- SEM, 2-way ANOVA with Sidak’s multiple comparisons test. F) Mean +/- SEM, 2-way ANOVA with Tukey’s multiple comparisons test. Non-significant comparisons are not indicated on the graph.

### CSPG4-targeting CAR-Ms alone do not cause reduced melanoma cell survival

We next determined whether CAR-M^αCSPG4^ phagocytosis affected melanoma cell survival. We first quantified the amount of melanoma cell trogocytosis since nibbling of target cells often does not result in cell death and compared this rate to whole cell phagocytosis. The median area of Lck-mScarlet melanoma cell fragments internalized in CAR-M^αCSPG4^ cells is significantly lower (65 μm^2^) than the median area of a whole, unengulfed Lck-mScarlet melanoma cell (473 μm^2^) (Supplemental Fig. 3E), suggesting that we were predominantly imaging trogocytosis events and not whole cell phagocytosis events. We next took advantage of image-based flow cytometry, where we could both mask for complete internalization, and quantify a relative size of the internalized fragment across thousands of cells (Fig. 3A, B). Based on the engulfment size measurements (Supplemental Fig. 3E), we stratified phagocytic events as CAR-Ms with large melanoma fragments (> 75 μm^2^) versus CAR-Ms with small melanoma fragments (< 75 μm^2^). We observed both a higher percent of overall phagocytosis, as well as a higher proportion of CAR-Ms with large melanoma fragments in CAR-M^αCSPG4^ compared to control CAR-M^GFP^ (Fig. 3C, D; Supplemental Fig. 4). To specifically quantify whole-cell phagocytosis, we engineered A375 cells to express a fluorescently-tagged histone H2B in the nucleus (H2B-mCherry), allowing us to measure nuclear engulfment as a read-out for whole-cell phagocytosis. Consistent with the image-based flow cytometry observations, this analysis revealed increased levels of whole-cell phagocytosis by CAR-M^αCSPG4^ versus CAR-M^αCSGP4ΔFcRγ^ or CAR-M^GFP^ controls (Fig. 3E). However, the overall level of whole-cell phagocytosis was low, at less than 2%. We then analyzed melanoma cell survival as the proportion of melanoma cells remaining in the coculture population and observed no significant changes in the overall percentage of melanoma cells in the coculture population across all conditions (Fig. 3F). These results suggest that although CAR-M^αCSPG4^ exhibited higher rates of whole cell melanoma phagocytosis compared to control CAR-Ms, these phagocytic events were not sufficient to reduce melanoma cell survival.

**Figure 3.**
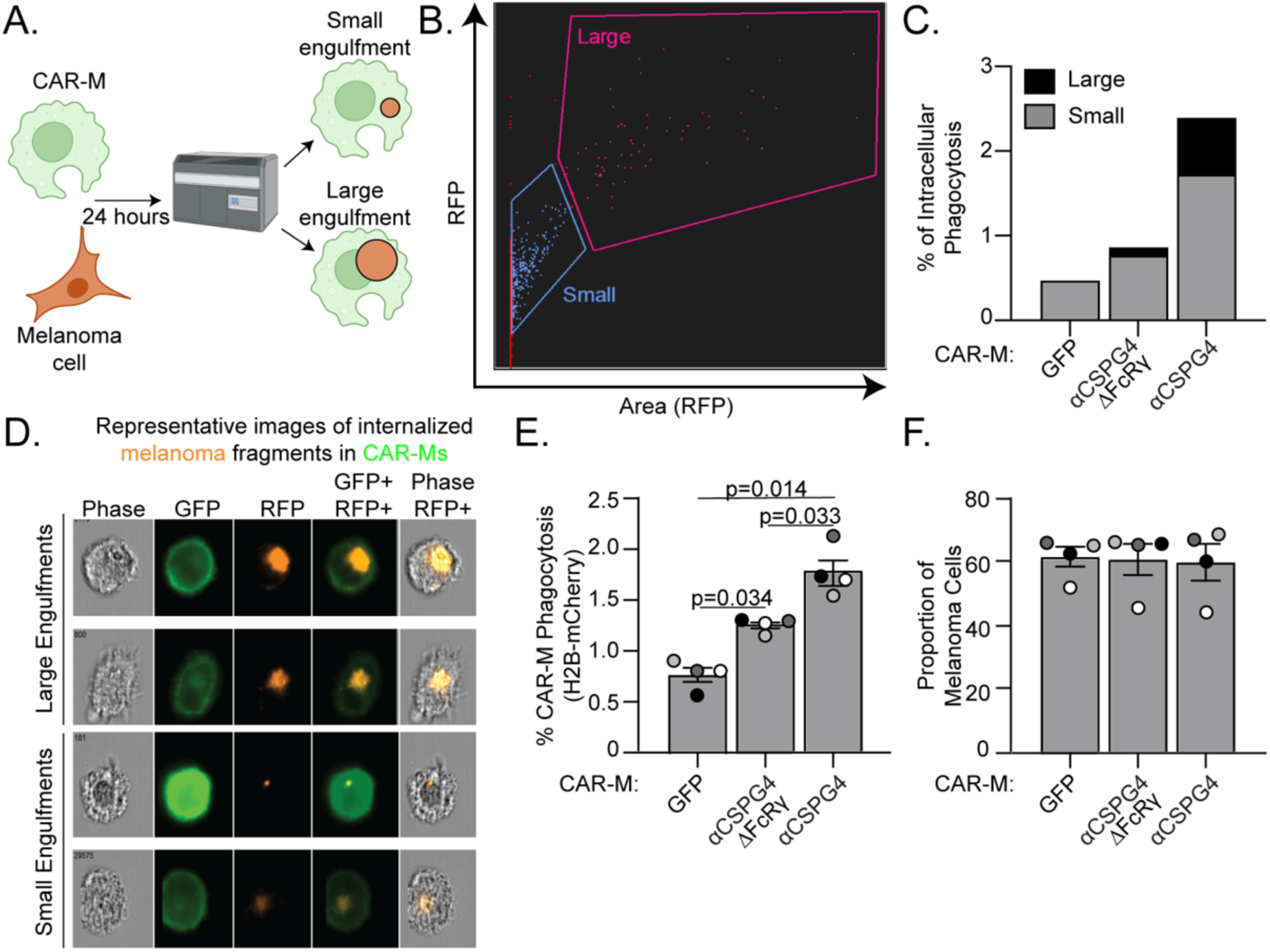
CSPG4-targeting CAR-Ms primarily trogocytose target melanoma cells. **A)** Schematic of CAR-M coculture with melanoma cells and analysis of bite size by image-based flow cytometry (Imagestream, Cytek Amnis). **B)** Representative gating of the size of internalized A375-Lck-mScarlet fragments in CAR-Ms, images acquired with a 63X objective lens on the Imagestream. **C)** Quantification of data in (B) – small (< 75 μm^2^) versus large (> 75 μm^2^) A375-Lck-mScarlet fragments inside CAR-Ms. n=1 biological donor. **D)** Representative images of “small” and “large” A375-Lck-mScarlet fragments (red) inside CAR-Ms (green). **E)** Quantification of percent CAR-M phagocytosis of A375-H2B-mCherry cells. **F)** Quantification of proportion of live, A375-H2B-mCherry cells remaining in the population by flow cytometry after 24 hours of coculture with CAR-M^αCSPG4^ compared to CAR-M^GFP^. Each dot on the graph is a biological replicate, each as a shade of gray. E, F) mean +/- SEM, 1-way ANOVA with Tukey’s multiple comparison test. Non-significant comparisons are not indicated on the graph.

### Combining CSPG4-targeting CAR-Ms with αCD47 inhibits melanoma growth in 3D

Puzzled by the lack of change in melanoma cell survival in our system, we next sought to test the effects of CAR-Ms in a more physiologically relevant 3D environment as macrophages in suspension compared to those adhered on tissue culture dishes exhibit a much greater capacity for phagocytosis (48). Many cancer cells, including melanoma, overexpress CD47 on their surface, a “don’t eat me” signal (49). Melanoma expressed CD47 interacts with SIRPα on macrophages, protecting the melanoma cells from phagocytosis (49). Blocking CD47 has been shown to inhibit A375 tumor growth in mice (50). Therefore, we hypothesized that elevated expression of CD47 in A375 cells inhibits CAR-M^αCSPG4^-mediated whole cell phagocytosis. We first cultured CAR-M/melanoma spheroids (Fig. 4A) using A375 cells expressing Lck-mScarlet and quantified phagocytosis with image-based flow cytometry after 24 hours. In 3D cultures with CAR-M^αCSPG4^, we observed a high level of melanoma phagocytosis to almost 50% at 24 hours, with a higher proportion of CAR-Ms with large, internalized melanoma fragments when Magrolimab (Hu5F9-G4), a humanized IgG4 monoclonal antibody that blocks the CD47/SIRPα axis (51–53), was added to the cultures compared to IgG controls (Supplemental Fig. 5A). Control CAR-M^GFP^ did not show altered phagocytosis of melanoma cells in the presence of αCD47 (Supplemental Fig. 5A). These results suggest that in 3D cultures, CAR-M^αCSPG4^ and αCD47 work cooperatively to enable phagocytosis. To test between trogocytosis and whole-cell phagocytosis, we again used melanoma cells with fluorescently tagged nuclei (H2B-mCherry). We observed a significant increase in the percent of CAR-M-mediated whole cell phagocytosis by CAR-M^αCSPG4^ compared to control CAR-M^GFP^, and that this whole cell phagocytosis was further increased to ∼35% with αCD47 (Fig. 4B). Excitingly, we found that the proportion of melanoma cells in the final spheroid was significantly reduced in the presence of CAR-M^αCSPG4^ with αCD47 (Fig. 4C). To determine whether the lack of melanoma cells was due to CAR-M-mediated melanoma cell death, we performed the 3D spheroid experiments and then dissociated the spheroids to visualize the cells with high-resolution microscopy. Dead cells exhibit ruptured and fragmented nuclei and histones become separated from DNA during apoptosis (54); thus, we quantified the number of internalized melanoma cells with ruptured H2B-mCherry nuclei as a measure of cell death. We observed a population of A375-H2B-mCherry cells that were not phagocytosed by CAR-Ms (“unengulfed”); a population of A375-H2B-mCherry cells that were engulfed by CAR-Ms, but the nuclei remained intact with the H2B-mCherry signal colocalizing with DAPI (“engulfed+live”); and a population of A375-H2B-mCherry cells that were engulfed by CAR-Ms with their nuclei broken down and H2B-mCherry no longer localized with DNA, indicated cell death (“engulfed+dead”). We found that a significantly higher proportion of A375 cells exhibited the “engulfed+dead” phenotype in cultures with CAR-M^αCSPG4^ and αCD47, compared to all other conditions (Fig. 4D, E; Supplemental Fig. 5B-D).

**Figure 4.**
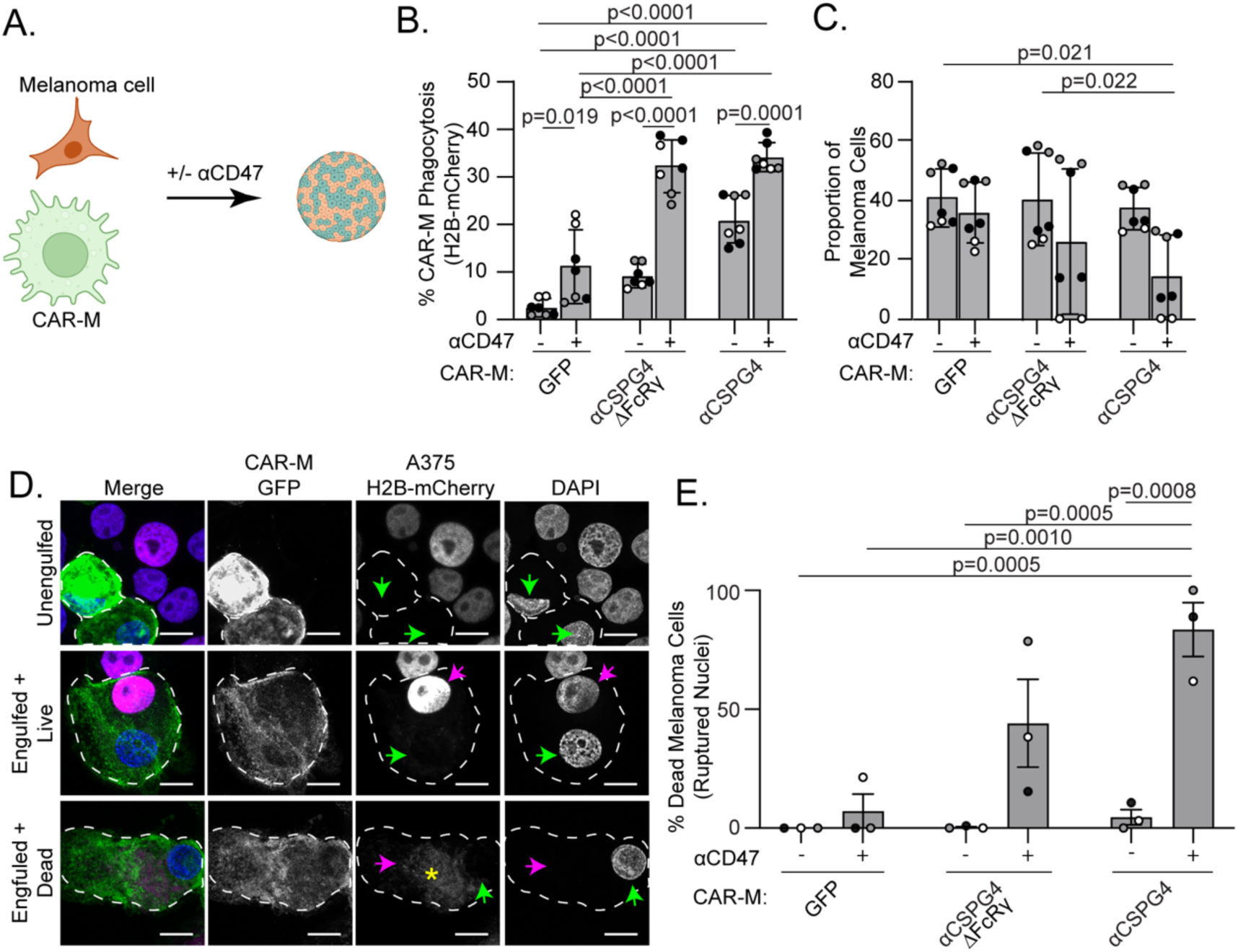
Combining CSPG4-targeting CAR-Ms with αCD47 leads to melanoma cell death in 3D. **A)** Schematic of CAR-M and melanoma cells co-forming 3D spheroids with and without αCD47. **B)** Quantification of percent CAR-M phagocytosis of melanoma cells in spheroids after 72 hours. **C)** Proportion of remaining melanoma cells in spheroids after 72 hours by flow cytometry. **D)** Representative images of replated spheroids showing “unengulfed” melanoma cells (single spherical nuclei, colocalization with H2B-mCherry, no engulfment by CAR-M); “engulfed + live” (melanoma engulfment by CAR-M, A375-H2B-mCherry nuclei intact and colocalizing with DAPI); or “engulfed + dead” (melanoma engulfment by CAR-M, A375-H2B-mCherry does not colocalize with DAPI). White dashed outline marks individual CAR-Ms. Green arrows indicate CAR-M nuclei. Magenta arrows indicate the A375-H2B-mCherry signal colocalizing with DAPI inside CAR-Ms. The asterisk indicates dispersed A375-H2B-mCherry signal not colocalizing with DAPI inside CAR-Ms. Scale bar indicates 10 microns. **E)** Quantification of ruptured nuclei as percent of total melanoma cells after 72 hours of coculture. CAR-M^GFP^/IgG (N=104), CAR-M^GFP^/αCD47 (N=70), CAR-M^αCSPG4ΔFcRγ^/IgG (N=110), CAR-M^αCSPG4ΔFcRγ^/αCD47 (N=47), CAR-M^αCSPG4^/IgG (N=65), CAR-M^αCSPG4^/ αCD47 (N=34). B, C, E) n=3 biological replicates; each shade of gray indicates one biological replicate, with each dot indicating technical replicates. Mean +/- SEM, 2-way ANOVA with Tukey’s multiple comparisons test. Non-significant comparisons are not indicated on the graphs.

Interestingly, the reduction in melanoma cells and robust CAR-M phagocytosis with αCD47 was only observed in 3D culture conditions and not observed in 2D cultures (Supplemental Fig. 5E-I). These results suggest that the combination of CAR-M^αCSPG4^ with αCD47 in 3D leads to high levels of melanoma cell death.

To better mimic tumor growth conditions, we altered the 3D approaches and allowed melanoma spheroids to form prior to addition of CAR-Ms (Fig. 5A). After 4-8 hours of CAR-M addition, we removed the spheroids from the U-bottom low-attachment plates and embedded them in Matrigel to visualize CAR-M interactions with the melanoma spheroid. We found that CAR-M^αCSPG4^ remain adhered to the melanoma spheroids when spheroids were transferred to Matrigel, whereas the CAR-M^GFP^ did not (Fig. 5B, C). Adding αCD47 to the culture did not affect CAR-M attachment to the spheroid. Using high resolution imaging, we further observed that CAR-M^αCSPG4^ infiltrate the spheroid (Supplemental Fig. 6A; Supplemental Video 3) and phagocytose melanoma cells within the spheroid (Supplemental Fig. 6B).

**Figure 5.**
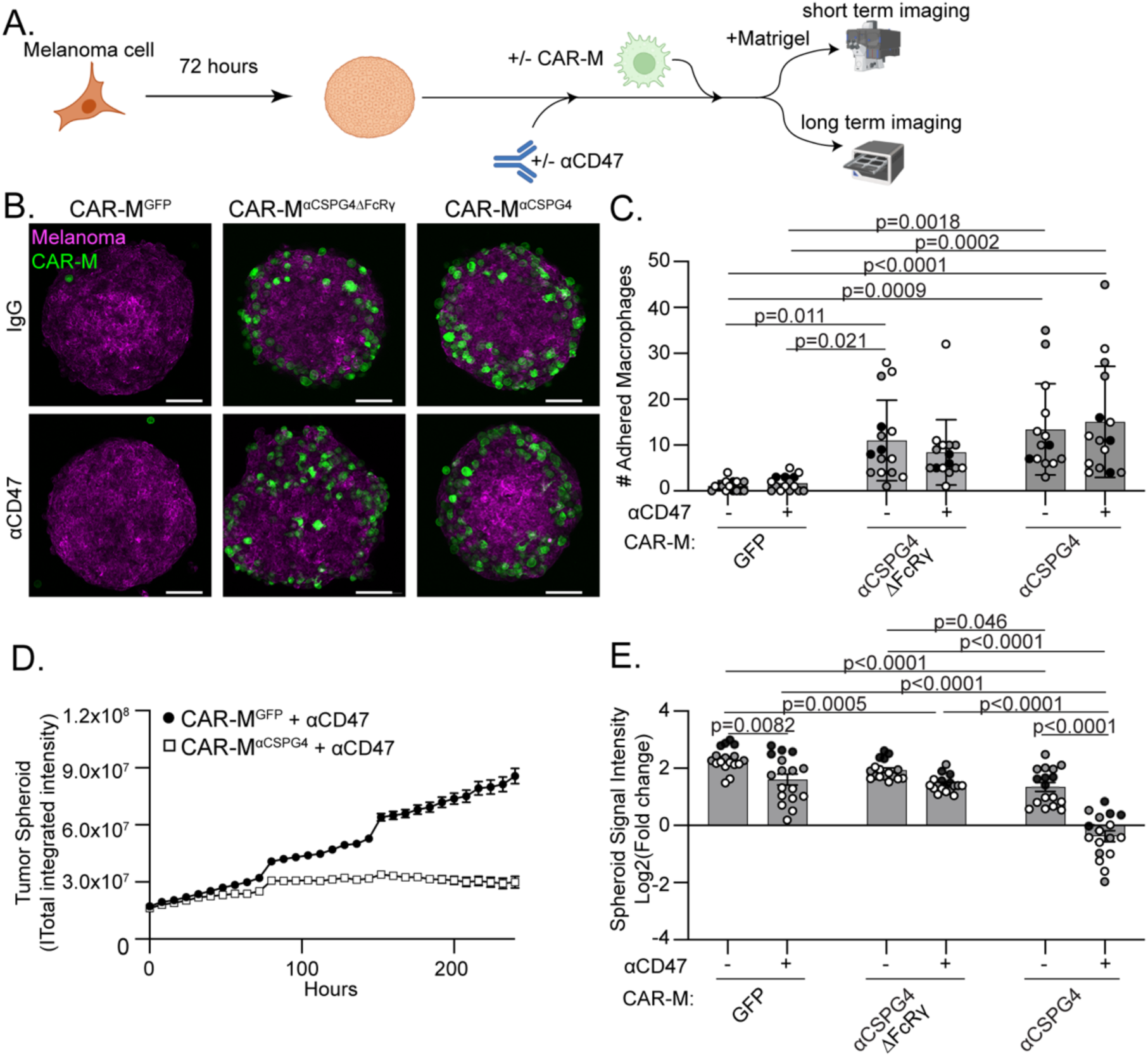
Combining CSPG4-targeting CAR-Ms with αCD47 inhibits melanoma spheroid growth in 3D. **A)** Schematic of pre-formed A375 melanoma spheroid, +/- αCD47, and CAR-Ms for short-term adherence assay and long-term imaging. **B)** Representative images of pre-formed A375- Lck-mScarlet spheroids (magenta) cultured with CAR-Ms (green) (following schematic in (A)) for 4-8 hours with αCD47 or IgG controls and embedded in Matrigel for imaging. Scale bar is 100 microns. **C)** Quantification of GFP+ CAR-Ms adhered to mScarlet+ melanoma spheroids in (B). **D)** Representative plot of A375-H2B-mCherry spheroid growth in culture with CSPG4-targeting CAR-Ms or control CAR-Ms over 10 days, as measured by the total integrated intensity of mCherry signal over time. **E)** Quantification of data in D) at day 10 across conditions (N = 17 spheroids across N=3 biological replicates). Data is shown as log_2_(fold change from time zero). For graphs, each shade of gray indicates one biological replicate, with each dot indicating technical replicates. C, E) Mean +/- SEM, 2-way ANOVA with Tukey’s multiple comparisons test. Non-significant comparisons are not indicated on the graph.

We next tested how CAR-Ms affect melanoma spheroid growth over time. We grew melanoma spheroids for 72 hours, added CAR-Ms, and then allowed spheroids to continue to grow for 10 days in the U-well low-attachment plates (Fig. 5A). We monitored spheroid growth over time and observed a synergistic effect in which combining CAR-M^αCSPG4^ with αCD47 resulted in robust inhibition of tumor spheroid growth (Fig. 5D, E; Supplemental Fig. 7A, B). Notably, CAR-M^αCSPG4^ infiltrated the melanoma spheroid; whereas, CAR-M^GFP^ localized primarily to the exterior on the melanoma spheroid (Supplemental Fig. 7C, D; Supplemental video 4), suggesting increased interactions between spheroids and CSPG4-targeting CAR-Ms versus controls. We next sought to determine whether this inhibition of spheroid growth was specific for cells expressing CSPG4; however, consistent with published literature, the A375 cells expressing the shCSPG4 clones failed to form viable spheroids and disassemble during media changes, thus precluding analysis (Supplemental Fig. 7E, F)(55). Taken together, these results suggest that combining CSPG4-targeting CAR-Ms with αCD47 approaches result in robust inhibition of melanoma growth in 3D by increased phagocytosis.

The dramatic synergistic effects of combining CSPG4-targeting CAR-Ms with αCD47 led us to question whether we could reduce the concentration of αCD47, and still observe a robust CAR-M-mediated phagocytosis. Reducing the αCD47 concentration, while still showing efficient CAR-M-mediated phagocytosis of cancer cells, may reduce off-target effects when used in patients given that almost all healthy cells express some level of CD47. We started with 10 µg/mL of αCD47 as used in previous work (53,56) and performed a serial dilution series. Combining αCD47 with control CAR-M^GFP^ showed no significant increase in melanoma phagocytosis until 10 µg/mL of αCD47 (Supplemental Fig. 7G; white bar at 10^4^ compared to IgG control). At this αCD47 concentration, we also observed a significant increase in CAR-M^αCSPG4^-mediated phagocytosis compared to CAR-M^GFP^ control (Supplemental Fig. 7G; black bar vs white bar at 10^4^). This level of CAR-M^αCSPG4^-mediated phagocytosis did not decrease until the concentration of αCD47 was reduced to 100 ng/mL αCD47 (Supplemental Fig. 7G; comparing black bars at 10^4^ to 10^2^). These results suggest that significantly decreased concentrations of αCD47 still enhance CSPG4-CAR-M-mediated phagocytosis of melanoma cells, but not with CAR-M^GFP^.

### CSPG4-targeting CAR-Ms inhibit melanoma growth *in vivo*

Given our *in vitro* findings, we sought to use a syngeneic system using murine melanoma cells and murine bone marrow-derived macrophages. We measured expression of CSPG4 in several mouse melanoma cell lines by flow cytometry and observed high CSPG4 expression (Supplemental Fig. 8A) in the YUMM1.7 and YUMM1.1 cell lines (57). We therefore generated YUMM1.7 cells expressing H2B-mCherry. We then determined which CSPG4 scFv to use for these murine studies. The 763.74 CSPG4 scFV has no demonstrated reactivity with the murine CSPG4 homolog, likely due to a two amino acid difference between human and mouse CSPG4 in the binding epitope (Supplemental Fig. 8B). Conversely, the 225.28 binding epitope only has a one amino acid difference between human and mouse CSPG4 and has previously demonstrated some cross-species reactivity (44). Thus, we first tested whether CAR-Ms generated with the 225.28 scFv phagocytosed murine YUMM1.7-H2B-mCherry melanoma cells in 3D (Supplemental Fig 8C). CAR-M^αCSPG4(225.28)^ exhibited a low rate of phagocytosis, with only ∼4% of CAR-M^αCSPG4(225.28)^ phagocytosing YUMM1.7-H2B-mCherry cells (Supplemental Fig. 8C), a rate similarly observed in CAR-M^αCSPG4(763.74)^ and control CAR-M^GFP^ cells. These results suggest that CAR-M^αCSPG4(225.28)^ do not effectively target murine CSPG4, and thus precluded our ability to test the function of CAR-M^αCSPG4(225.28)^ in syngeneic mouse models. We therefore turned our attention to testing human CAR-Ms and human melanoma cells in immune-compromised models.

We determined whether CAR-M^αCSPG4(763.74)^ inhibits A375 human melanoma growth in immune-compromised NRG mice (58). We opted to inject CAR-Ms peri-tumorally, rather than systemically (11), to maximize on-target effects, while minimizing potential off-target effects in other tissues associated with intravenous administration. We first determined how long CAR-Ms are detectable in the tumor to determine the treatment regimen. We reasoned that the CAR-Ms did not need to stay in the tumor for the entirety of the experiment, but CAR-Ms must be present in the tumor long enough to phagocytose CSPG4-positive melanoma cells and potentially reprogram the tumor environment for a sustained response. Thus, we injected 400K A375 cells into the right flank of 6-8 week-old female NRG mice, waited for tumors to become palpable, and then peri-tumorally injected 1M CAR-Ms on day 9. We then euthanized mice on days 12, 14, 16, and 19 to determine whether CAR-Ms were present in the tumor by flow cytometry (Fig. 6A). We found that CAR-Ms were present at 1% of the total live cells in the tumor after 3 days post-injection (day 12), and that this number dropped to almost zero starting at 5 days post-injection (day 14) (Fig. 6B) Thus, we proceeded with a two-stage CAR-M injection strategy, 3-5 days apart.

**Figure 6.**
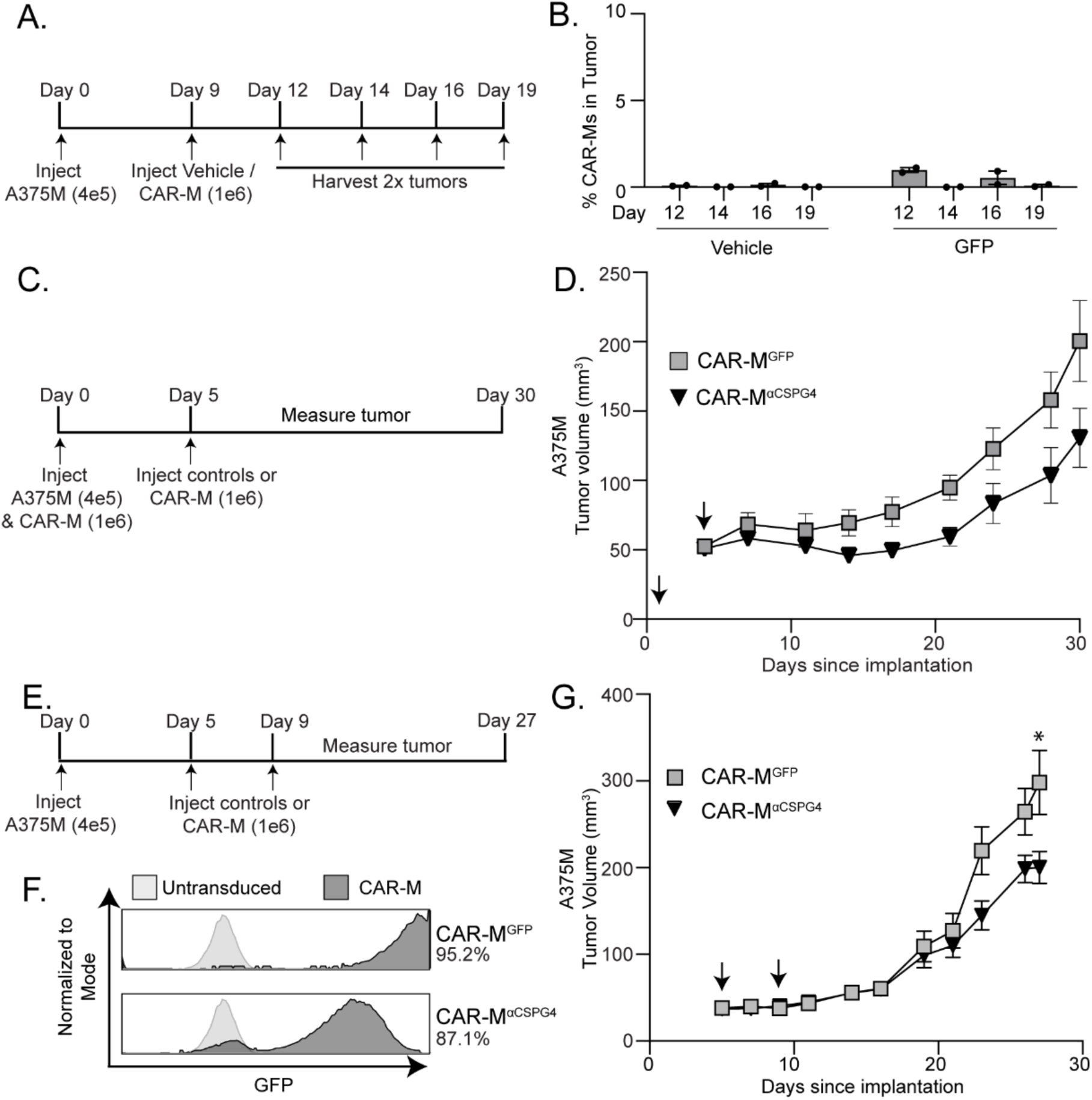
αCSPG4 CAR-Ms inhibit melanoma growth *in vivo*. **A)** Experimental outline showing that A375-H2B-mCherry cells were injected in NRG mice, and then injected with CAR-M^GFP^ after 9 days. Tumors were dissociated and processed for flow cytometry on indicated days to determine presence of CAR-Ms. **B)** Quantification of the presence of CAR-M^GFP^ in tumors over time in (A). **D)** Experimental outline showing A375-H2B-mCherry same-day injections with CAR-M^αCSPG4^ or CAR-M^GFP^ in NRG mice (N=4 mice per group). **D)** Tumor volume for (C) plotted over time. Arrows indicate CAR-M injections. **E)** Experimental outline showing CAR-M^αCSPG4^ or CAR-M^GFP^ injections after A375-H2B-mCherry engraftment in NRG mice (N=6 mice per group). **F)** Flow cytometry plots showing percent bone marrow-derived macrophage transduction for CAR-M^GFP^ and CAR-M^αCSPG4^. **G)** Tumor volume for (E, F) plotted over time. Arrows indicate CAR-M injections. D, G) 1-way ANOVA with Tukey’s multiple comparisons test at terminal time point. * p < 0.05 versus indicated group.

We first tested subcutaneous same-day injections of A375-H2B-mCherry cells and CAR-Ms into the flanks of mice, as same-day injection approaches were routinely used in previously published CAR-M work (11,13,14,17). We injected 400K A375-H2B-mCherry cells and 1M CAR-M^αCSPG4^, CAR-M^GFP^, or vehicle control and then re-injected with only CAR-Ms or vehicle control again 5 days later (Fig. 6C). We found no differences in animal weight across conditions, suggesting no gross off-target toxicity (Supplemental Fig. 9A). We found that treating mice with CAR-M^αCSPG4^ inhibited the growth of the tumor compared to CAR-M^GFP^ controls, although these differences were not statistically significant (Fig. 6D, Supplemental Fig. 9B-C). As expected, treatment with control CAR-M^GFP^ also inhibited melanoma growth compared to vehicle controls (Supplemental Fig. 9B, C), suggesting that the addition of any exogenous macrophages alone affects tumor growth as has been previously described (11,14,17). Together these results suggest that CSPG4-targeting CAR-Ms may reduce melanoma growth *in vivo*.

We next modeled CAR-M treatment following tumor growth, and thus moved beyond same-day injection studies. We injected A375-H2B-mCherry cells into NRG mice, and first allowed the tumors to engraft for 5 days (Fig. 6E). We then peritumorally injected CAR-M^αCSPG4^ or controls (Fig. 6F) on day 5 and again on day 9. We again found no differences in animal weight across conditions (Supplemental Fig. 9D). Excitingly, we found that CAR-M^αCSPG4^-treated animals exhibited a significant reduction in tumor growth compared to CAR-M^GFP^-treated animals (Fig. 6G; Supplemental Fig. 9E, F), and that this inhibition of tumor growth began at day 22 and was maintained through day 27, well after CAR-Ms are no longer detected in the tumor.

Immunofluorescence of the tumors at day 27 confirmed that majority of CAR-Ms are no longer present at the tumor (Supplemental Fig. 9G), suggesting that the sustained inhibition of melanoma growth is likely mediated by reprogramming of the myeloid cells present in the tumor microenvironment. These results show that first, CSPG4-targeting CAR-Ms effectively inhibit melanoma growth *in vivo*, and second, the inhibition of tumor growth occurs in the absence of adaptive immune cells, suggesting the innate immune system is sufficient for this response.

## DISCUSSION

Current therapies for treating advanced stages of melanoma have improved patient survival, but primary resistance is still a relatively common occurrence, as well as recurrence via acquired resistance (27). Pro-tumorigenic macrophages constitute a large portion of the tumor microenvironment in melanoma. Since CAR-T cells have been unsuccessful at infiltrating solid tumors, we hypothesized that CAR-Ms may effectively target melanoma tumors. In this study, we unveiled a potential treatment strategy for melanoma by CSPG4-targeting CAR-Ms. Our findings demonstrate that CSPG4-targeting CAR-Ms efficiently phagocytose metastatic melanoma cells. When CSPG4-targeting CAR-Ms are combined with CD47 blocking antibodies, thereby blocking a key tumor cell “don’t eat me” signal, we observe enhanced phagocytosis of melanoma cells and robust inhibition of melanoma spheroid growth in 3D. Importantly, we show that CSPG4-targeting CAR-Ms control tumor growth *in vivo*. We demonstrate that in a xenograft model, with mice lacking adaptive immune cells (58), CSPG4-targeting CAR-Ms exhibit sustained inhibition of melanoma growth. These data support a growing body of work presenting CSPG4 as a potential therapeutic target, and that CAR-Ms may be effective at clearing cancer cells that overexpress CSPG4, particularly when combined with αCD47.

Our findings raise important questions regarding how long CAR-Ms must persist at the tumor to maintain tumor control. If CAR-M-mediated phagocytosis of cancer cells is the primary mechanism for anti-tumor activity, then both sustained persistence and delivery of high CAR-M numbers is critical. Alternatively, if tumor immune reprogramming by CAR-Ms is the primary mechanism for anti-tumor activity, transient CAR-M persistence at the tumor may be sufficient for tumor control. Both mechanisms likely contribute to anti-tumor activity, and further work is required to parse out the contributions of each of these mechanisms. Our data show that CAR-Ms are not detectable in the melanoma tumor after approximately 5 days. This short time frame is consistent with other reports (11,16). Despite the short-term CAR-M persistence in the tumor, as in this study, CAR-Ms targeting multiple tumor antigens across a variety of models maintain tumor control (11,13,24). These results and ours suggest that CAR-M-mediated phagocytosis of cancer cells is not sufficient to prevent tumor growth alone, and likely requires contributions from other stromal cells. We provide evidence that CSPG4-targeting CAR-Ms inhibit melanoma growth in immune-compromised animals over 18 days from the last CAR-M injection, well after CAR-Ms are no longer detected in the tumor (58). These results suggest that innate immune cells are sufficient to inhibit melanoma growth in animals treated with CSPG4-targeted CAR-Ms, likely by local reprogramming of the tumor-associated macrophages into anti-tumor macrophages, thereby increasing phagocytosis of melanoma cells by endogenous macrophages, and/or by decreasing immunosuppressive signals at the tumor and promoting general anti-tumor responses. We expect that while we show that innate immune cells inhibit melanoma growth, CAR-Ms likely also regulate the adaptive immune response, as has been previously suggested (11,13,22,59). Future CAR-M studies must perform systematic immune profiling at multiple time points over the growth of the tumor, dissecting each immune cell subset, their activation patterns, and exhaustion markers to provide a more in-depth understanding of the dynamics driving immune cell reprogramming in the tumor during CAR-M therapies. Furthermore, the number of CAR-Ms required, the injection frequency, as well as whether local versus systemic CAR-M injections provide better anti-tumor responses need to be systematically tested and compared.

Teasing apart the requirement for CAR-M persistence at the tumor and CAR-M-mediated immune reprogramming will guide the development of next-generation CAR-Ms and combination therapies. We hypothesized that increasing CAR-M-mediated phagocytosis of cancer cells is a critical aspect of anti-tumor activity for three reasons: 1) increase cancer cell cytotoxicity directly through phagocytosis; 2) facilitate the repolarization of tumor-associated macrophages into anti-tumor macrophages; 3) increase antigen presentation to activate the adaptive immune arm. Thus, we focused our efforts on improving CAR-M-mediated phagocytosis of melanoma cells by engineering the CSPG4-targeting chimeric antigen receptor to contain the intracellular FcRγ phagocytic signaling domain and combining the CSPG4-targeting CAR-Ms with CD47 blocking antibodies. It remains to be seen whether this combination will prove effective for melanoma *in vivo*, however other solid tumor models have demonstrated success with pairing αCD47 strategies with CAR-Ms both *in vitro* and *in vivo* (12,24,25,60). Future approaches to improve phagocytosis include using activation domains more specific to the engulfment machinery such as the Rac2E62K mutation (19) and targeting additional “don’t eat me” signals (61). If sustained CAR-M persistence at the tumor is deemed to be critical for tumor growth control, then efforts to improve CAR-M survival such as overexpression of cytokines like M-CSF may be needed (62). Another approach to improve CAR-M phagocytosis activity is to directly polarize CAR-Ms to a pro-inflammatory phenotype prior to administration (11). This approach would have the benefit of immediately releasing cytokines to dampen the immunosuppressive environment of the tumor; however, it remains unknown whether this approach increases the potential for dangerous side-effects such as cytokine release syndrome (11). Alternative approaches include combining CAR-Ms with ICB treatments. Due to the antigen presentation capabilities of macrophages, the strength of CAR-Ms may lie in their ability to phagocytose cancer cell targets and display additional tumor antigens to T-cells as a mechanism to overcome tumor heterogeneity. Combining CSPG4-CAR-Ms with ICB approaches may represent a powerful approach to tackle the long-standing challenge of tumor heterogeneity particularly in melanoma (63). Preliminary results suggest that combining CAR-Ms with ICB therapies can have an additive effect in reducing tumor burden, however if it is unclear how this combination affects tumor immune populations or ICB resistance (60). Our work provides compelling evidence that CSPG4-targeting CAR-Ms successfully reduces primary melanoma growth across *in vitro* and *in vivo* melanoma models. The next critical steps involve determining whether local administration of CSPG4-targeting CAR-Ms at the time of melanoma tumor resection prevents local melanoma recurrence. Furthermore, future work is required to determine whether systemic versus local administration of CSPG4-targeting CAR-Ms prevent melanoma metastases.

## ACKNOWLEDGMENTS

We would like to acknowledge Dr. Meghan Morrisey and members of the Roh-Johnson lab for constructive feedback and advice; and Kate Modzelewska for outstanding technical support. This work was supported by National Cancer Institute R37CA247994, the Huntsman Cancer Institute Translational Science Initiative Award 24-TSI-02, and the University of Utah Ascender Grant U-8006 (to MRJ); S.R. Lowry Endowed Chair in the Department of Neurosurgery (to SHC); Neurosurgery Research Education Foundation Grant 10067440 (to EK); Department of Defense Melanoma Research Program W81XWH2210495 (to RLB); National Cancer Institute of the National Institutes of Health P30CA042014 (to RLJ-T); Huntsman Cancer Institute Cancer Center Support Grant P30CA040214, the American Cancer Society IRG-21-131-01, and Five For The Fight (to MQR); Office Of The Director of the National Institutes of Health S10OD026959 and National Cancer Institute 5P30CA042014-24 (to Flow Cytometry Core); and National Cancer Institute P30CA042014 (to Preclinical Shared Resource).

## AUTHOR CONTRIBUTIONS

DG: conceptualization, investigation, data curation, writing – original draft, writing – review & editing; QX: investigation, data curation; TW: investigation, data curation; EK: investigation, data curation; RLB: data curation, investigation; RLJ-T: investigation, resources, funding acquisition; MQR: data curation, investigation; SHC & GD: resources; MRJ: conceptualization, investigation, data curation, resources, writing – original draft, writing – review & editing, supervision, funding acquisition.

## DATA AVAILABILITY STATEMENT

The data generated in this study are available within the article and its corresponding supplementary data files. Expression profile data analyzed in this study were obtained from Gene Expression Omnibus (GEO) at GSE151091(36), the [Broad Single Cell portal](https://singlecell.broadinstitute.org/single_cell/study/SCP109/melanoma-immunotherapy-resistance)(37), and the human protein atlas(64).

## REFERENCES

1. June CH, Sadelain M. Chimeric Antigen Receptor Therapy. N Engl J Med 2018;379(1):64–73 doi 10.1056/NEJMra1706169.

2. Anurathapan U, Chan RC, Hindi HF, Mucharla R, Bajgain P, Hayes BC, et al. Kinetics of tumor destruction by chimeric antigen receptor-modified T cells. Mol Ther 2014;22(3):623–33 doi 10.1038/mt.2013.262.

3. Kuczek DE, Larsen AMH, Thorseth ML, Carretta M, Kalvisa A, Siersbæk MS, et al. Collagen density regulates the activity of tumor-infiltrating T cells. J Immunother Cancer 2019;7(1):68 doi 10.1186/s40425-019-0556-6.

4. Gerber AL, Munst A, Schlapbach C, Shafighi M, Kiermeir D, Husler R, et al. High expression of FOXP3 in primary melanoma is associated with tumour progression. Br J Dermatol 2014;170(1):103–9 doi 10.1111/bjd.12641.

5. Salmi S, Siiskonen H, Sironen R, Tyynelä-Korhonen K, Hirschovits-Gerz B, Valkonen M, et al. The number and localization of CD68+ and CD163+ macrophages in different stages of cutaneous melanoma. Melanoma Res 2019;29(3):237–47 doi 10.1097/CMR.0000000000000522.

6. Mirlekar B. Tumor promoting roles of IL-10, TGF-beta, IL-4, and IL-35: Its implications in cancer immunotherapy. SAGE Open Med 2022;10:20503121211069012 doi 10.1177/20503121211069012.

7. Noman MZ, Hasmim M, Messai Y, Terry S, Kieda C, Janji B, et al. Hypoxia: a key player in antitumor immune response. A Review in the Theme: Cellular Responses to Hypoxia. Am J Physiol Cell Physiol 2015;309(9):C569–79 doi 10.1152/ajpcell.00207.2015.

8. Fu Z, Mowday AM, Smaill JB, Hermans IF, Patterson AV. Tumour Hypoxia-Mediated Immunosuppression: Mechanisms and Therapeutic Approaches to Improve Cancer Immunotherapy. Cells 2021;10(5) doi 10.3390/cells10051006.

9. Herbel C, Patsoukis N, Bardhan K, Seth P, Weaver JD, Boussiotis VA. Clinical significance of T cell metabolic reprogramming in cancer. Clin Transl Med 2016;5(1):29 doi 10.1186/s40169-016-0110-9.

10. Noy R, Pollard JW. Tumor-associated macrophages: from mechanisms to therapy. Immunity 2014;41(1):49–61 doi 10.1016/j.immuni.2014.06.010.

11. Klichinsky M, Ruella M, Shestova O, Lu XM, Best A, Zeeman M, et al. Human chimeric antigen receptor macrophages for cancer immunotherapy. Nat Biotechnol 2020;38(8):947–53 doi 10.1038/s41587-020-0462-y.

12. Morrissey MA, Williamson AP, Steinbach AM, Roberts EW, Kern N, Headley MB, et al. Chimeric antigen receptors that trigger phagocytosis. Elife 2018;7 doi 10.7554/eLife.36688.

13. Lei A, Yu H, Lu S, Lu H, Ding X, Tan T, et al. A second-generation M1-polarized CAR macrophage with antitumor efficacy. Nat Immunol 2024;25(1):102–16 doi 10.1038/s41590-023-01687-8.

14. Zhang L, Tian L, Dai X, Yu H, Wang J, Lei A, et al. Pluripotent stem cell-derived CAR-macrophage cells with antigen-dependent anti-cancer cell functions. J Hematol Oncol 2020;13(1):153 doi 10.1186/s13045-020-00983-2.

15. Dong X, Fan J, Xie W, Wu X, Wei J, He Z, et al. Efficacy evaluation of chimeric antigen receptor-modified human peritoneal macrophages in the treatment of gastric cancer. Br J Cancer 2023;129(3):551–62 doi 10.1038/s41416-023-02319-6.

16. Zhang W, Liu L, Su H, Liu Q, Shen J, Dai H, et al. Chimeric antigen receptor macrophage therapy for breast tumours mediated by targeting the tumour extracellular matrix. Br J Cancer 2019;121(10):837–45 doi 10.1038/s41416-019-0578-3.

17. Zhang J, Webster S, Duffin B, Bernstein MN, Steill J, Swanson S, et al. Generation of anti-GD2 CAR macrophages from human pluripotent stem cells for cancer immunotherapies. Stem Cell Reports 2023;18(2):585–96 doi 10.1016/j.stemcr.2022.12.012.

18. Chuang ST, Stein JB, Nevins S, Kilic Bektas C, Choi HK, Ko WK, et al. Enhancing CAR Macrophage Efferocytosis Via Surface Engineered Lipid Nanoparticles Targeting LXR Signaling. Adv Mater 2024:e2308377 doi 10.1002/adma.202308377.

19. Mishra AK, Rodriguez M, Torres AY, Smith M, Rodriguez A, Bond A, et al. Hyperactive Rac stimulates cannibalism of living target cells and enhances CAR-M-mediated cancer cell killing. Proc Natl Acad Sci U S A 2023;120(52):e2310221120 doi 10.1073/pnas.2310221120.

20. Liu M, Liu J, Liang Z, Dai K, Gan J, Wang Q, et al. CAR-Macrophages and CAR-T Cells Synergistically Kill Tumor Cells In Vitro. Cells 2022;11(22) doi 10.3390/cells11223692.

21. Kang SW, You S, Wong EA, El Halawani ME. Normalization of transfection efficiency using the beta-lactamase gene of the pGL3 luciferase vector in primary anterior pituitary cells. Biotechniques 2002;33(2):326–8, 30 doi 10.2144/02332st05.

22. Niu Z, Chen G, Chang W, Sun P, Luo Z, Zhang H, et al. Chimeric antigen receptor-modified macrophages trigger systemic anti-tumour immunity. J Pathol 2021;253(3):247–57 doi 10.1002/path.5585.

23. Huo Y, Zhang H, Sa L, Zheng W, He Y, Lyu H, et al. M1 polarization enhances the antitumor activity of chimeric antigen receptor macrophages in solid tumors. J Transl Med 2023;21(1):225 doi 10.1186/s12967-023-04061-2.

24. Chen C, Jing W, Chen Y, Wang G, Abdalla M, Gao L, et al. Intracavity generation of glioma stem cell-specific CAR macrophages primes locoregional immunity for postoperative glioblastoma therapy. Sci Transl Med 2022;14(656):eabn1128 doi 10.1126/scitranslmed.abn1128.

25. Chen Y, Zhu X, Liu H, Wang C, Chen Y, Wang H, et al. The application of HER2 and CD47 CAR-macrophage in ovarian cancer. J Transl Med 2023;21(1):654 doi 10.1186/s12967-023-04479-8.

26. Program SR. 2024 April 17. SEER*Explorer: An interactive website for SEER cancer statistics. National Cancer Institute <seer.cancer.gov/statistics-network/explorer/>. Accessed 2024 April 17.

27. Carlino MS, Larkin J, Long GV. Immune checkpoint inhibitors in melanoma. Lancet 2021;398(10304):1002–14 doi 10.1016/S0140-6736(21)01206-X.

28. Wilson BS, Imai K, Natali PG, Ferrone S. Distribution and molecular characterization of a cell-surface and a cytoplasmic antigen detectable in human melanoma cells with monoclonal antibodies. Int J Cancer 1981;28(3):293–300 doi 10.1002/ijc.2910280307.

29. Price MA, Colvin Wanshura LE, Yang J, Carlson J, Xiang B, Li G, et al. CSPG4, a potential therapeutic target, facilitates malignant progression of melanoma. Pigment Cell Melanoma Res 2011;24(6):1148–57 doi 10.1111/j.1755-148X.2011.00929.x.

30. Geldres C, Savoldo B, Hoyos V, Caruana I, Zhang M, Yvon E, et al. T lymphocytes redirected against the chondroitin sulfate proteoglycan-4 control the growth of multiple solid tumors both in vitro and in vivo. Clin Cancer Res 2014;20(4):962–71 doi 10.1158/1078-0432.CCR-13-2218.

31. Center ULCC, Pharmaceuticals B, Center UCRFaLCC. Autologous CAR-T Cells Targeting CSPG4 in Relapsed/Refractory HNSCC. https://classic.clinicaltrials.gov/show/NCT06096038; 2024.

32. Landoni E, Fuca G, Wang J, Chirasani VR, Yao Z, Dukhovlinova E, et al. Modifications to the Framework Regions Eliminate Chimeric Antigen Receptor Tonic Signaling. Cancer Immunol Res 2021;9(4):441–53 doi 10.1158/2326-6066.CIR-20-0451.

33. Greiner D, Scott TM, Olson GS, Aderem A, Roh-Johnson M, Johnson JS. Genetic Modification of Primary Human Myeloid Cells to Study Cell Migration, Activation, and Organelle Dynamics. Curr Protoc 2022;2(8):e514 doi 10.1002/cpz1.514.

34. Weischenfeldt J, Porse B. Bone Marrow-Derived Macrophages (BMM): Isolation and Applications. CSH Protoc 2008;2008:pdb.prot5080 doi 10.1101/pdb.prot5080.

35. Schindelin J, Arganda-Carreras I, Frise E, Kaynig V, Longair M, Pietzsch T, et al. Fiji: an open-source plasorm for biological-image analysis. Nat Methods 2012;9(7):676–82 doi 10.1038/nmeth.2019.

36. Belote RL, Le D, Maynard A, Lang UE, Sinclair A, Lohman BK, et al. Human melanocyte development and melanoma dedifferentiation at single-cell resolution. Nat Cell Biol 2021;23(9):1035–47 doi 10.1038/s41556-021-00740-8.

37. Jerby-Arnon L, Shah P, Cuoco MS, Rodman C, Su MJ, Melms JC, et al. A Cancer Cell Program Promotes T Cell Exclusion and Resistance to Checkpoint Blockade. Cell 2018;175(4):984–97 e24 doi 10.1016/j.cell.2018.09.006.

38. Riaz N, Havel JJ, Makarov V, Desrichard A, Urba WJ, Sims JS, et al. Tumor and Microenvironment Evolution during Immunotherapy with Nivolumab. Cell 2017;171(4):934–49.e16 doi 10.1016/j.cell.2017.09.028.

39. Yang J, Zhao S, Wang J, Sheng Q, Liu Q, Shyr Y. A pan-cancer immunogenomic atlas for immune checkpoint blockade immunotherapy. Cancer Res 2021;82(4):539–42 doi 10.1158/0008-5472.CAN-21-2335.

40. Ilieva KM, Cheung A, Mele S, Chiarutni G, Crescioli S, Griffin M, et al. Chondroitin Sulfate Proteoglycan 4 and Its Potential As an Antibody Immunotherapy Target across Different Tumor Types. Front Immunol 2017;8:1911 doi 10.3389/fimmu.2017.01911.

41. Tirosh I, Izar B, Prakadan SM, Wadsworth MH, 2nd, Treacy D, Trombetta JJ, et al. Dissecting the multicellular ecosystem of metastatic melanoma by single-cell RNA-seq. Science 2016;352(6282):189-96 doi 10.1126/science.aad0501.

42. Beard RE, Zheng Z, Lagisetty KH, Burns WR, Tran E, Hewitt SM, et al. Multiple chimeric antigen receptors successfully target chondroitin sulfate proteoglycan 4 in several different cancer histologies and cancer stem cells. J Immunother Cancer 2014;2:25 doi 10.1186/2051-1426-2-25.

43. Cheng CY, Mruk DD. The blood-testis barrier and its implications for male contraception. Pharmacol Rev 2012;64(1):16–64 doi 10.1124/pr.110.002790.

44. Williams IP, Crescioli S, Sow HS, Bax HJ, Hobbs C, Ilieva KM, et al. In vivo safety profile of a CSPG4-directed IgE antibody in an immunocompetent rat model. MAbs 2020;12(1):1685349 doi 10.1080/19420862.2019.1685349.

45. Mittelman A, Tiwari R, Lucchese G, Willers J, Dummer R, Kanduc D. Identification of monoclonal anti-HMW-MAA antibody linear peptide epitope by proteomic database mining. J Invest Dermatol 2004;123(4):670–5 doi 10.1111/j.0022-202X.2004.23417.x.

46. Temponi M, Gold AM, Ferrone S. Binding parameters and idiotypic profile of the whole immunoglobulin and Fab’ fragments of murine monoclonal antibody to distinct determinants of the human high molecular weight-melanoma associated antigen. Cancer Res 1992;52(9):2497–503.

47. Geiser M, Schultz D, Le Cardinal A, Voshol H, García-Echeverría C. Identification of the human melanoma-associated chondroitin sulfate proteoglycan antigen epitope recognized by the antitumor monoclonal antibody 763.74 from a peptide phage library. Cancer Res 1999;59(4):905–10.

48. Cannon GJ, Swanson JA. The macrophage capacity for phagocytosis. J Cell Sci 1992;101 ( Pt 4):907–13 doi 10.1242/jcs.101.4.907.

49. Ngo M, Han A, Lakatos A, Sahoo D, Hachey SJ, Weiskopf K, et al. Antibody Therapy Targeting CD47 and CD271 Effectively Suppresses Melanoma Metastasis in Patient-Derived Xenograws. Cell Rep 2016;16(6):1701–16 doi 10.1016/j.celrep.2016.07.004.

50. Wang Y, Ni H, Zhou S, He K, Gao Y, Wu W, et al. Tumor-selective blockade of CD47 signaling with a CD47/PD-L1 bispecific antibody for enhanced anti-tumor activity and limited toxicity. Cancer Immunol Immunother 2021;70(2):365–76 doi 10.1007/s00262-020-02679-5.

51. Sikic BI, Lakhani N, Patnaik A, Shah SA, Chandana SR, Rasco D, et al. First-in-Human, First-in-Class Phase I Trial of the Anti-CD47 Antibody Hu5F9-G4 in Patients With Advanced Cancers. J Clin Oncol 2019;37(12):946–53 doi 10.1200/JCO.18.02018.

52. Willingham SB, Volkmer JP, Gentles AJ, Sahoo D, Dalerba P, Mitra SS, et al. The CD47-signal regulatory protein alpha (SIRPa) interaction is a therapeutic target for human solid tumors. Proc Natl Acad Sci U S A 2012;109(17):6662–7 doi 10.1073/pnas.1121623109.

53. Gholamin S, Mitra SS, Feroze AH, Liu J, Kahn SA, Zhang M, et al. Disrupting the CD47-SIRPα anti-phagocytic axis by a humanized anti-CD47 antibody is an efficacious treatment for malignant pediatric brain tumors. Sci Transl Med 2017;9(381) doi 10.1126/scitranslmed.aaf2968.

54. Wu D, Ingram A, Lahti JH, Mazza B, Grenet J, Kapoor A, et al. Apoptotic release of histones from nucleosomes. J Biol Chem 2002;277(14):12001–8 doi 10.1074/jbc.M109219200.

55. Uno K, Koya Y, Yoshihara M, Iyoshi S, Kitami K, Sugiyama M, et al. Chondroitin Sulfate Proteoglycan 4 Provides New Treatment Approach to Preventing Peritoneal Dissemination in Ovarian Cancer. Int J Mol Sci 2024;25(3) doi 10.3390/ijms25031626.

56. Zhang M, Hutter G, Kahn SA, Azad TD, Gholamin S, Xu CY, et al. Anti-CD47 Treatment Stimulates Phagocytosis of Glioblastoma by M1 and M2 Polarized Macrophages and Promotes M1 Polarized Macrophages In Vivo. PLoS One 2016;11(4):e0153550 doi 10.1371/journal.pone.0153550.

57. Meeth K, Wang JX, Micevic G, Damsky W, Bosenberg MW. The YUMM lines: a series of congenic mouse melanoma cell lines with defined genetic alterations. Pigment Cell Melanoma Res 2016;29(5):590–7 doi 10.1111/pcmr.12498.

58. Pearson T, Shultz LD, Miller D, King M, Laning J, Fodor W, et al. Non-obese diabetic-recombination activating gene-1 (NOD-Rag1 null) interleukin (IL)-2 receptor common gamma chain (IL2r gamma null) null mice: a radioresistant model for human lymphohaematopoietic engrawment. Clin Exp Immunol 2008;154(2):270–84 doi 10.1111/j.1365-2249.2008.03753.x.

59. Kang M, Lee SH, Kwon M, Byun J, Kim D, Kim C, et al. Nanocomplex-Mediated In Vivo Programming to Chimeric Antigen Receptor-M1 Macrophages for Cancer Therapy. Adv Mater 2021;33(43):e2103258 doi 10.1002/adma.202103258.

60. Wang X, Su S, Zhu Y, Cheng X, Cheng C, Chen L, et al. Metabolic Reprogramming via ACOD1 depletion enhances function of human induced pluripotent stem cell-derived CAR-macrophages in solid tumors. Nat Commun 2023;14(1):5778 doi 10.1038/s41467-023-41470-9.

61. Khalaji A, Yancheshmeh FB, Farham F, Khorram A, Sheshbolouki S, Zokaei M, et al. Don’t eat me/eat me signals as a novel strategy in cancer immunotherapy. Heliyon 2023;9(10):e20507 doi 10.1016/j.heliyon.2023.e20507.

62. Kim AB, Xiao Q, Yan P, Pan Q, Pandey G, Grathwohl S, et al. Chimeric antigen receptor macrophages target and resorb amyloid plaques. JCI Insight 2024;9(6) doi 10.1172/jci.insight.175015.

63. Ng MF, Simmons JL, Boyle GM. Heterogeneity in Melanoma. Cancers (Basel) 2022;14(12) doi 10.3390/cancers14123030.

64. Karlsson M, Zhang C, Mear L, Zhong W, Digre A, Katona B, et al. A single-cell type transcriptomics map of human tissues. Sci Adv 2021;7(31) doi 10.1126/sciadv.abh2169.

