## Supplemental Figures 1-9 for "CSPG4-targeting CAR-macrophages inhibit melanoma growth"

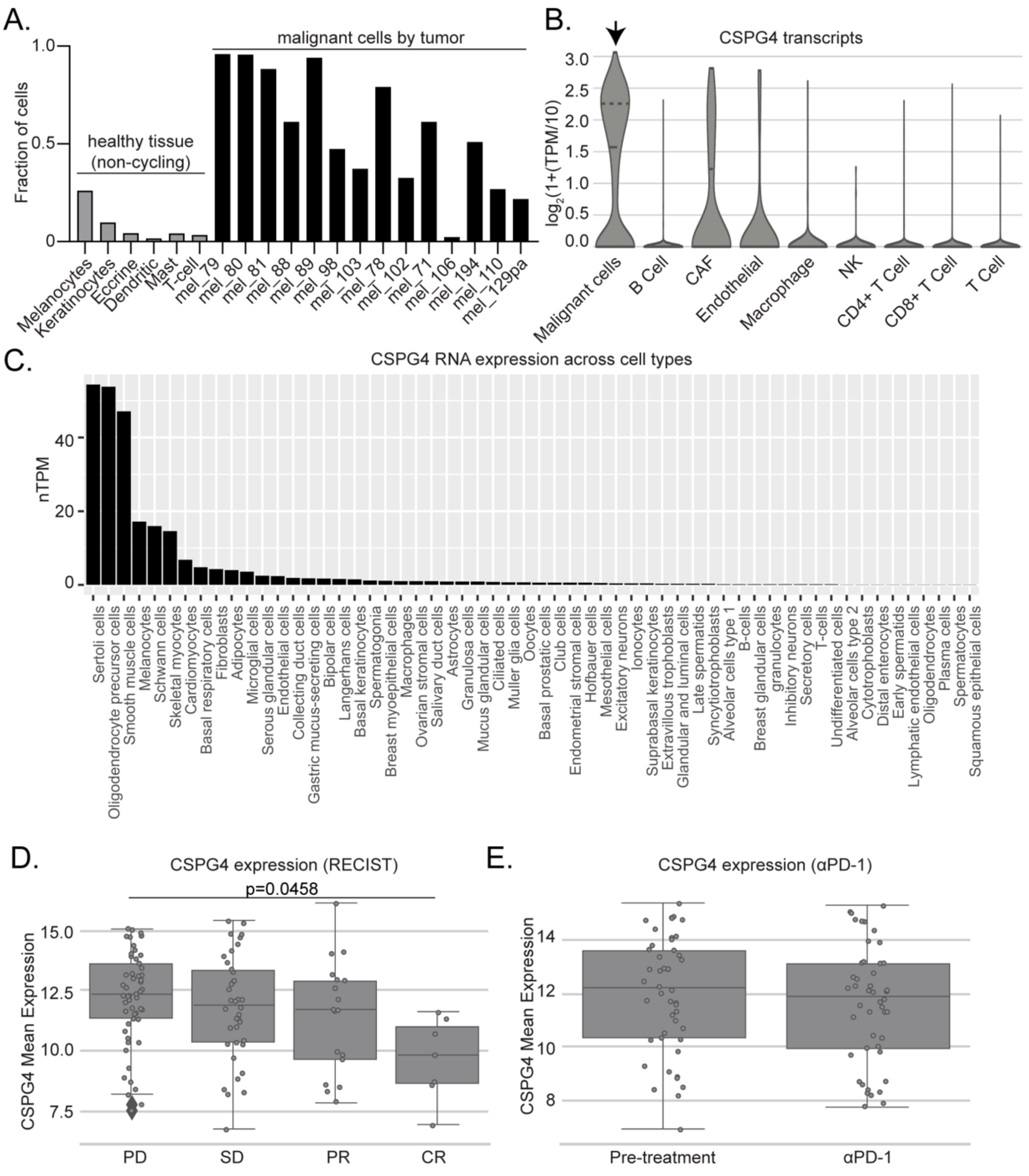


Supplemental Fig 1

**Supplemental Figure 1|** CSPG4 expression is overexpressed and specific to melanoma. **A)** Fraction of cells expressing CSPG4 in single-cell RNA sequencing analysis of healthy (non-cycling, gray) tissue vs melanoma tumors (black) rank mean normalized from Figure 1A. **B)** Violin plot of CSPG4 transcript levels by single-cell RNA sequencing analysis of malignant cells (arrow) in tumors compared to non-malignant cells within the tumor. Quartiles are demarcated on the violin plots. **C)** CSPG4 transcript counts in normal cell types (Human Protein Atlas). **D, E)** Box plots of CSPG4 expression from the Melanoma-Riaz study on the Immuno-genomic atlas for immune checkpoint blockade-based cancer therapy by RECIST status (D) (PD = Progressive disease, SD = stable disease, PR = partial response, CR = complete response), and by pre-treatment vs post-αPD-1/L1 treatment (E). D) 1-way ANOVA with Tukey's multiple comparisons test. E) Unpaired t-test, non-significant comparisons are not indicated on the graph.

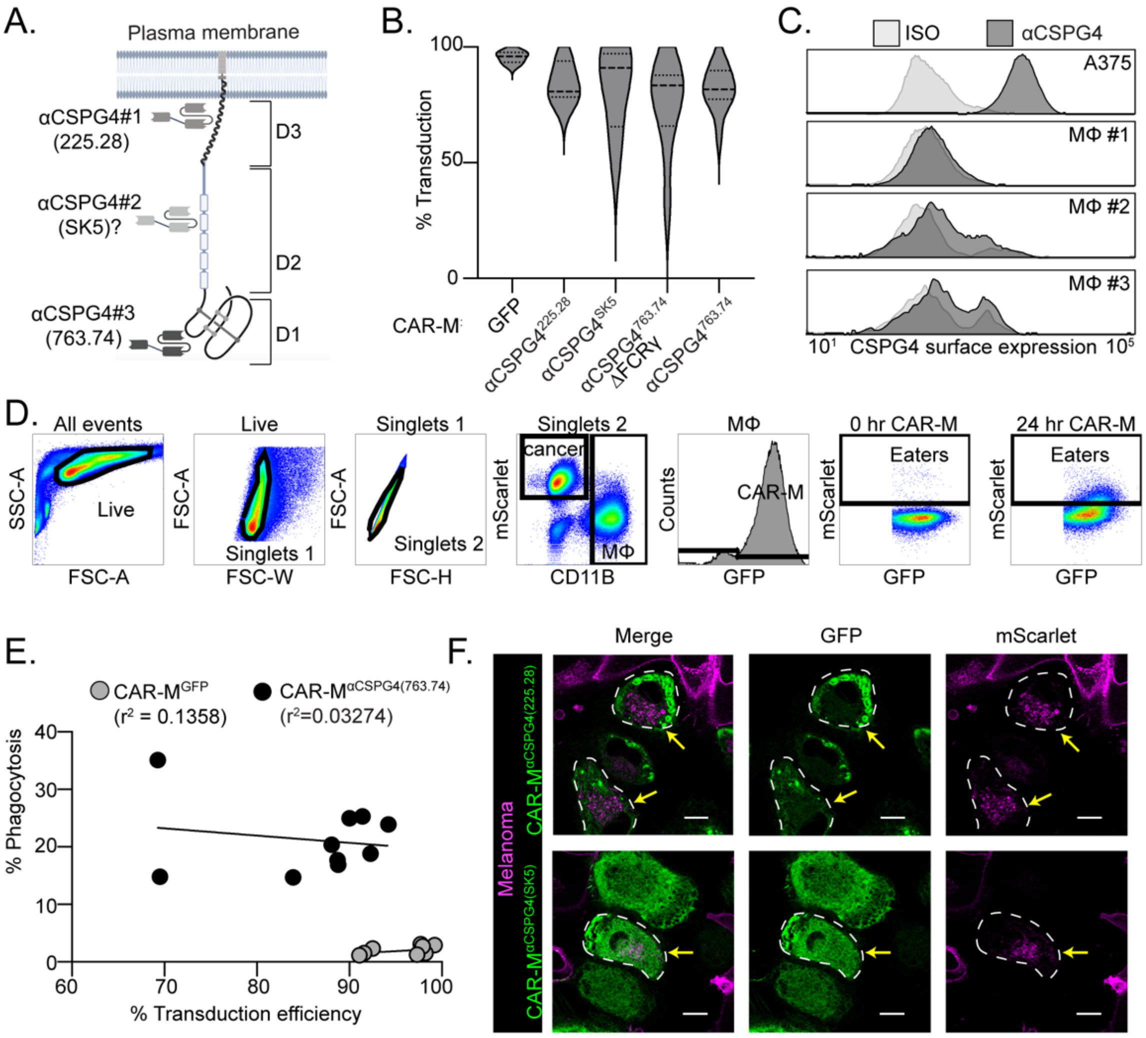


Supplemental Fig 2

**Supplemental Figure 2|** Phagocytosis gating strategy and CAR-M transduction efficiencies. **A)** Schematic of CSPG4 scFvs showing predicted (SK5) and validated (225.28, 763.74) domain binding. **B)** Violin plot of primary macrophage transduction efficiency for each CAR-M construct (N: GFP = 12, 763.74 = 12, SK5 = 4, 225.28 = 3). **C)** CSPG4 surface expression by flow cytometry of A375 cells compared to primary macrophages (n=3 donors as MΦ #1-3). **D)** Gating strategy for quantifying CAR-M transduction and CAR-M-mediated phagocytosis of A375-Lck-mScarlet cells based on 0-hour coculture. **E)** Comparison of CAR-M$^{\alpha CSPG4(763.74)}$ and CAR-M$^{GFP}$ phagocytosis and transduction efficiency from flow cytometry experiments with line of best fit (slope - CAR-M$^{GFP}$: 0.086 and CAR-M$^{\alpha CSPG4}$: -0.13) and correlation. **F)** Representative single z-plane images of CAR-M$^{\alpha CSPG4(SK5)}$ or CAR-M$^{\alpha CSPG4(225.28)}$ (green) and A375-Lck-mScarlet cells (magenta). The white dashed line outlines CAR-M$^{\alpha CSPG4}$ cell boundary, and the yellow arrows highlight engulfment events. Images acquired with a 63X objective on an LSM 880 microscope. Scale bar is 10 microns.

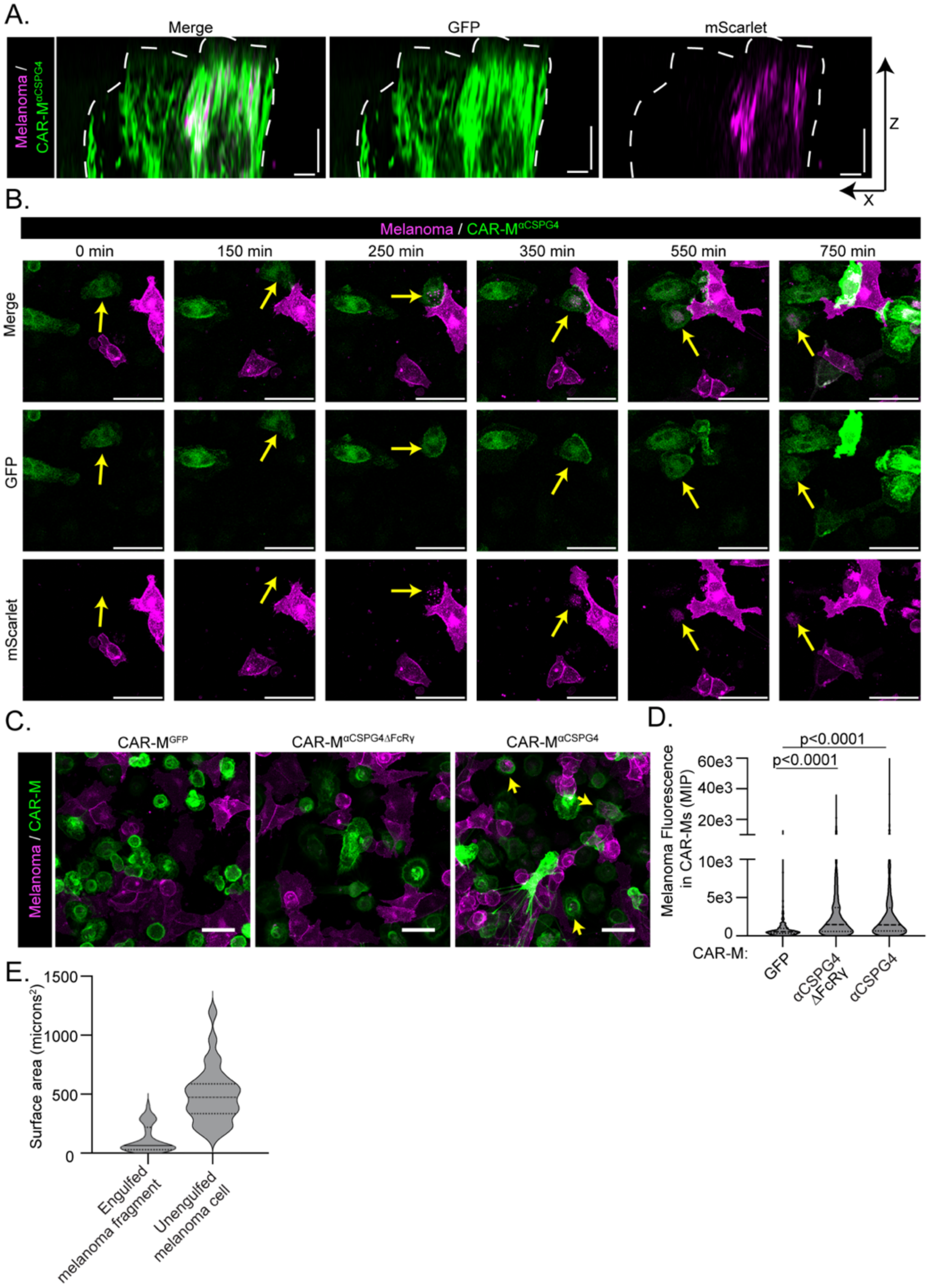


**Supplemental Figure 3|** Melanoma particles are fully internalized within CSPG4-targeting CAR-Ms. **A)** Representative X-Z plane images of engulfed Lck-mScarlet fragments inside of CAR-M$^{\alpha CSPG4}$ cells shown in Fig 1G. Images acquired with a 63X objective on an LSM 880 microscope. Scale bars on both x and z axis represent 5 microns. **B)** Representative still images from timelapse recording (Supplemental Video 2) of CAR-M$^{\alpha CSPG4}$ (green) engulfment of Lck-mScarlet fragments (magenta). Yellow arrows highlight engulfment events. Scale bar is 50 microns. **C)** Representative images of A375-Lck-mScarlet cells cocultured with CAR-M$^{\alpha CSPG4}$ or CAR-M$^{GFP}$ for 24 hours, images taken at 20X, yellow arrows indicate CAR-M phagocytosis, scale bar is 40 microns. **D)** Violin plot, with quartiles, of quantification of Lck-mScarlet signal overlapping with GFP+ CAR-Ms in (C) from a maximum intensity projection (n=3 biological replicates). Mean +/- SEM, 1-way ANOVA with Tukey's multiple comparisons test. **E)** Violin plot, with quartiles, of surface area measured for of internalized Lck-mScarlet fragments in CAR-M$^{\alpha CSPG4}$ (N= 34 engulfments, median = 64.69 microns$^2$) or unengulfed A375-Lck-mScarlet cells (N = 28 cells, median = 473.19 microns$^2$).

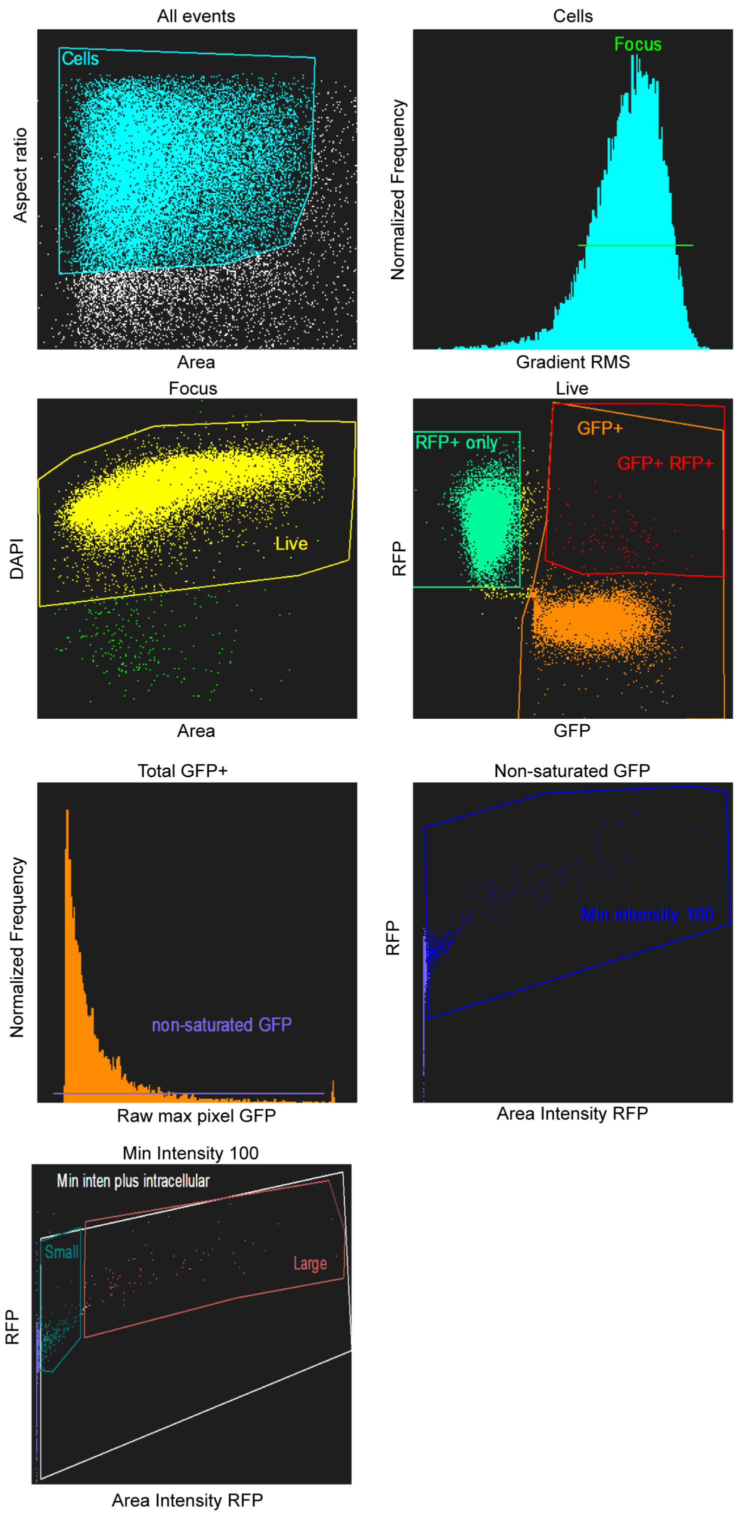


Supplemental Fig 4

**Supplemental Figure 4|** Imagestream gating strategy for phagocytic events. Imagestream gating strategy for identification of internalized Lck-mScarlet+ melanoma cell fragments in GFP+ CAR-M cells (data in Figure 3).

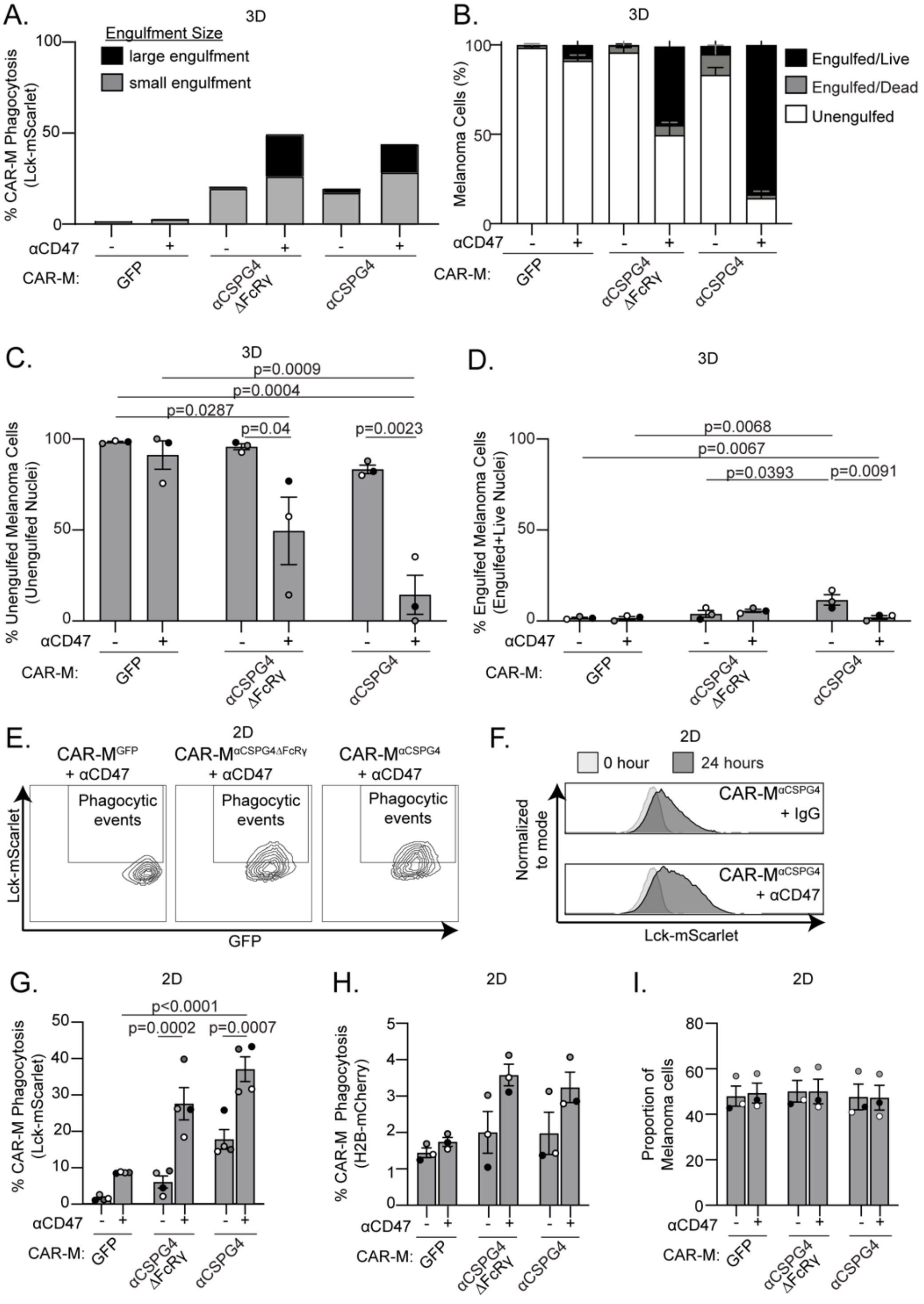


Supplemental Fig 5

**Supplemental Figure 5|** αCD47 treatment increases CSPG4-CAR-M-mediated melanoma cell death in 3D, but not in 2D. **A)** Imagestream quantification of fully internalized A375 Lck-mScarlet puncta in GFP+ CAR-M cells in 3D spheroids after 24 hours; images acquired with a 40X objective lens on an Imagestream. **B)** Classification of melanoma cells (live or dead) after 3 days of coculture with CAR-Ms as co-formed spheroids from Figure 4D. **C)** Quantification of unengulfed nuclei as percent of total melanoma cells after 72 hours of coculture in B. **D)** Quantification of engulfed nuclei as percent of total melanoma cells after 72 hours of coculture in B. **E)** Representative flow cytometry contour plots of CAR-M phagocytosis of A375-Lck-mScarlet cells treated with IgG isotype control or 10 μg/mL αCD47 for 24 hours in 2D culture. **F)** A normalized to mode histogram of Lck-mScarlet fluorescence in CAR-Ms at 0 and 24 hours in CAR-M$^{\alpha CSPG4}$ with and without αCD47 in 2D culture. **G)** Quantification of CAR-M phagocytosis of A375-Lck-mScarlet cells treated with and without αCD47 in (E, F). **H)** Quantification of CAR-Ms phagocytosis of A375-H2B-mCherry cells treated with IgG isotype control or 10 μg/mL αCD47 in 2D culture. **I)** Proportion of target A375-H2B-mCherry cells remaining after 24 hours of coculture with CAR-Ms from (H). Each dot is 1 biological replicate, as shades of gray. C, D, G, H, I) Mean +/- SEM, 2-way AVOVA with Tukey's multiple comparisons test. Non-significant comparisons are not indicated on the graphs.

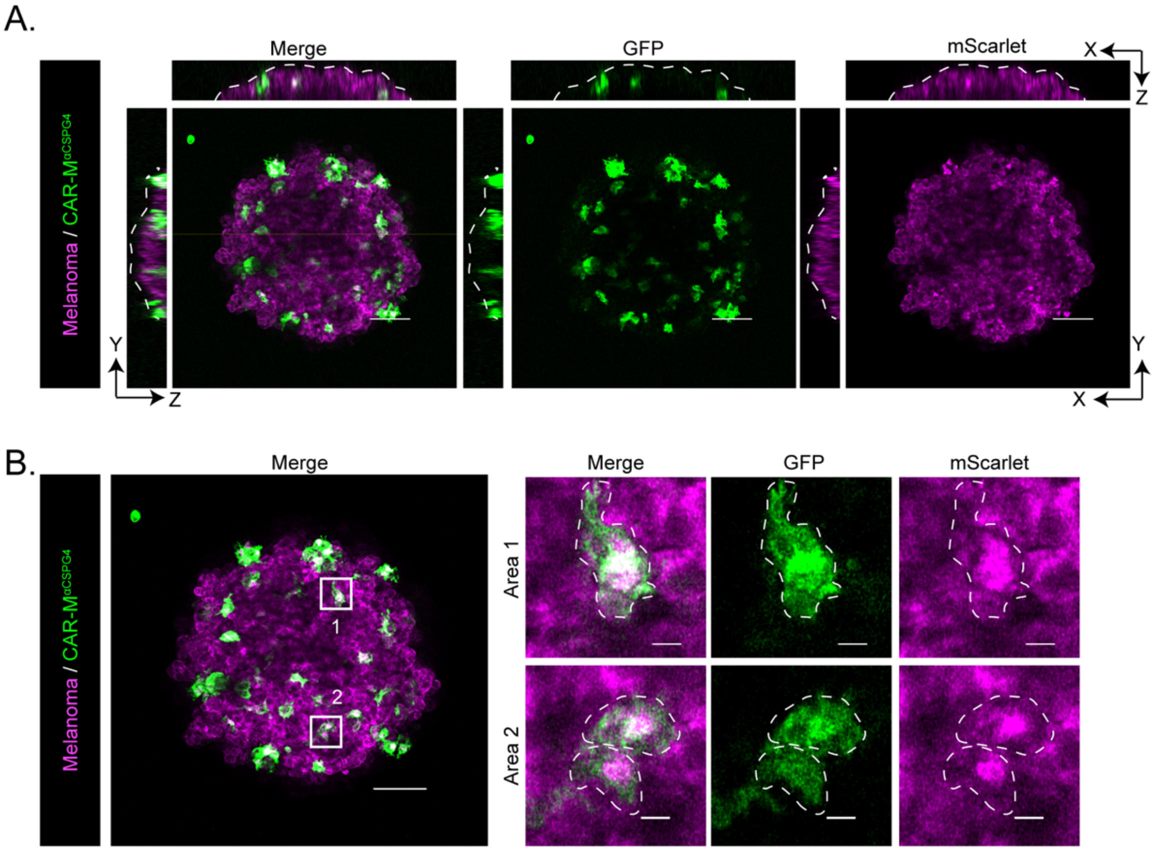


Supplemental Fig 6

**Supplemental Figure 6|** CSPG4 CAR-Ms infiltrate melanoma spheroids and phagocytose melanoma cells. **A)** Representative still images with orthogonal x-z, and y-z slices of CAR-M$^{\alpha CSPG4}$ (green) inside A375-Lck-mScarlet spheroids (magenta) after 72 hours of coculture. Spheroid boundary is marked by white dashed line. Scale bar is 100 microns. Images acquired with a 10X objective. **B)** Representative maximum intensity projection of 3 slices indicating CAR-M (green) engulfment of mScarlet+ fragments. Two areas of the spheroid (boxes) are magnified to the right as Area 1 and Area 2, with CAR-Ms outlined in white hatched lines. Scale bar of main image is 100 microns; scale bar of inset is 10 microns.

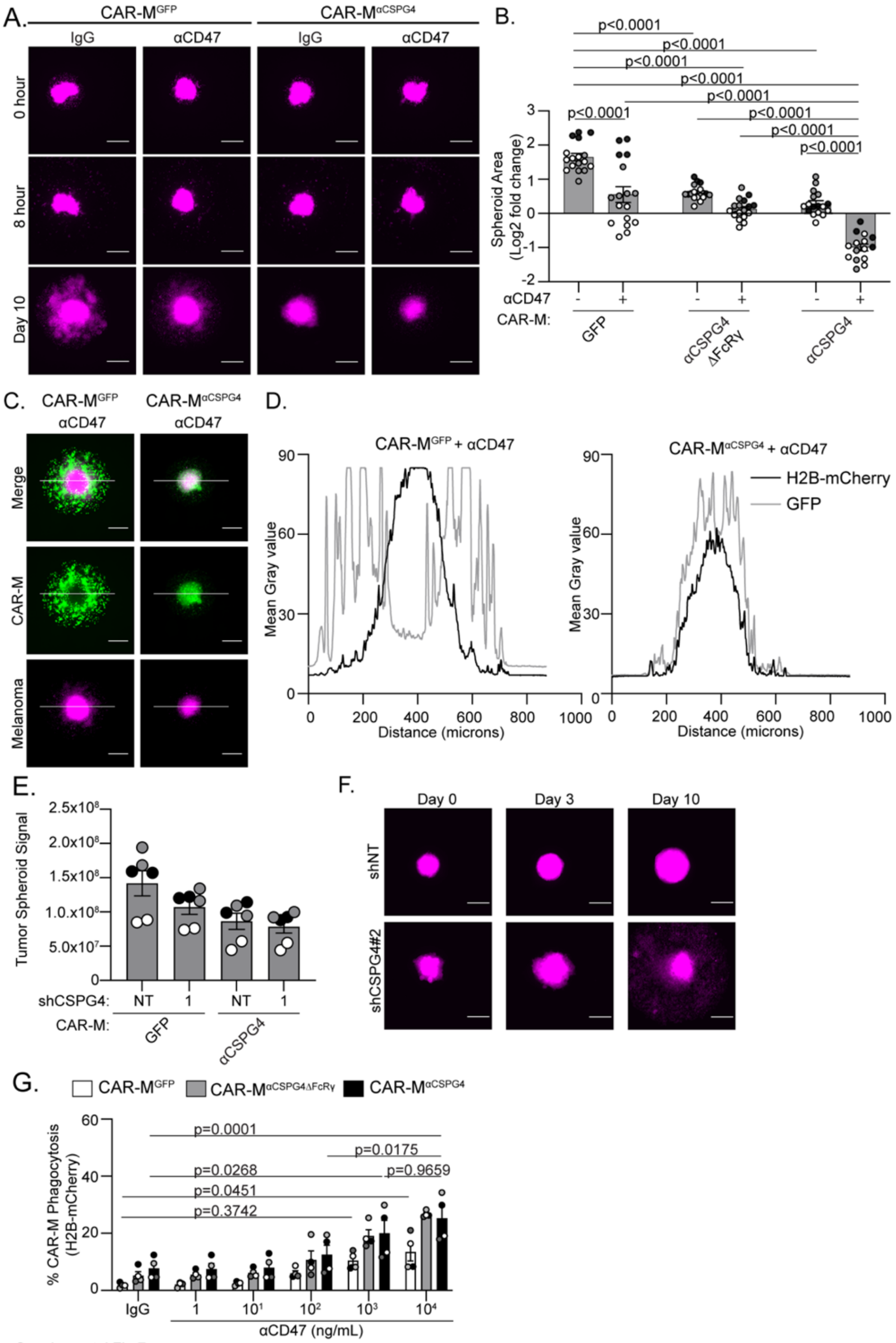


Supplemental Fig 7

**Supplemental Figure 7|** Combining CSPG4-targeting CAR-Ms with αCD47 approaches leads to whole-cell melanoma phagocytosis in 3D. **A)** Representative images of pre-formed A375-H2B-mCherry spheroids (magenta) at day 0 (before CAR-Ms), 8 hours after CAR-M addition, and 10 days after CAR-M addition. Scale bar is 200 microns. **B)** Quantification of A375 spheroid growth as measured by mCherry area at day 10 across conditions (N = 17 spheroids across N=3 biological replicates). Data shown as $log_2$(fold change from time zero). Mean +/- SEM, 2-way ANOVA with Dunnett's multiple comparisons test. **C)** Representative images of pre-formed A375-

H2B-mCherry spheroids (magenta) and CAR-Ms (green) on day 10; white line indicates line scan measurements in (D). Scale bar is 200 microns. **D)** Line scan signal intensity from representative spheroid images in C. **E)** Quantification of mScarlet integrated density in A375-Lck-mScarlet tumor spheroids after 10 days. A375-Lck-mScarlet cells were either treated with shNT or shCSPG4 RNA. **F)** Representative images of shNT and shCSPG4 #2 clones at day 0, day 3 (first media change), and day 10. Scale bar is 200 microns. **G)** Flow cytometry CAR-M phagocytosis quantification of A375-H2B-mCherry cells after 72 hours of 3D coculture with decreasing concentrations of αCD47. G) Mean +/- SEM, 2-way ANOVA with Tukey's multiple comparisons test. N=4 biological replicates. Non-significant comparisons are not indicated on the graphs, with the exception of on (G).

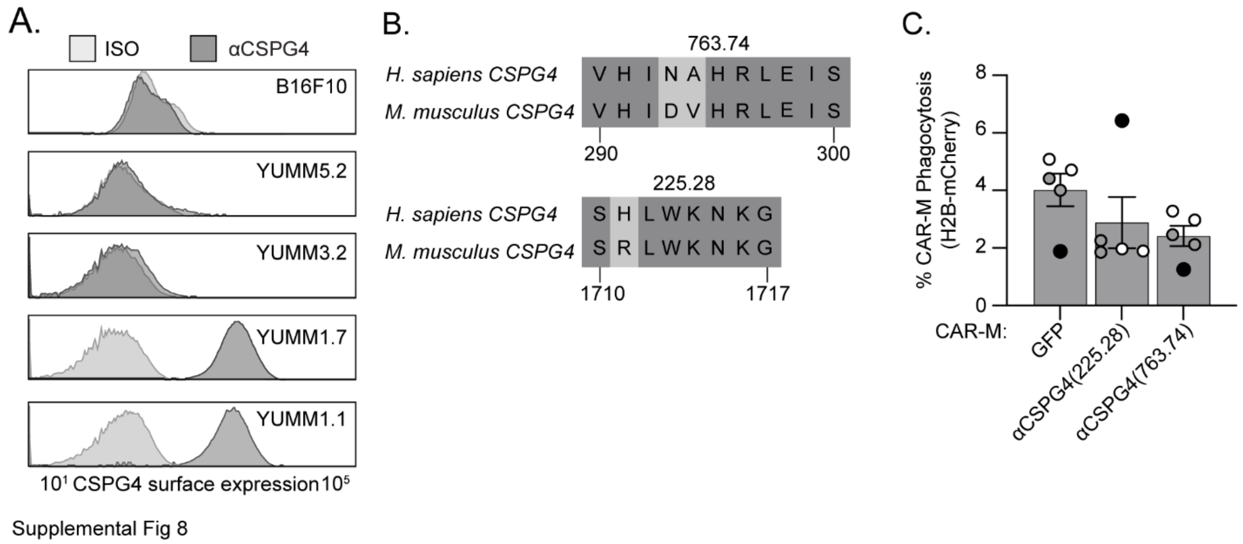


**Supplemental Figure 8|** CSPG4-targeting CAR-Ms using the 225.28 scFv do not efficiently phagocytose murine melanoma cells. **A)** Representative flow cytometry histograms of CSPG4 surface expression across a panel of murine melanoma cell lines. **B)** UniProt alignment of human vs mouse CSPG4 amino acid sequences for epitopes identified for 763.74 and 225.28 scFvs. **C)** Quantification of CSPG4$^{\alpha CSPG4(225.28)}$ phagocytosis of YUMM1.7 murine melanoma cells compared to CAR-M$^{GFP}$ and CAR-M$^{\alpha CSPG4(763.74)}$ by flow cytometry. 1-way ANOVA with Tukey's multiple comparisons test, non-significant comparisons are not indicated on the graph.

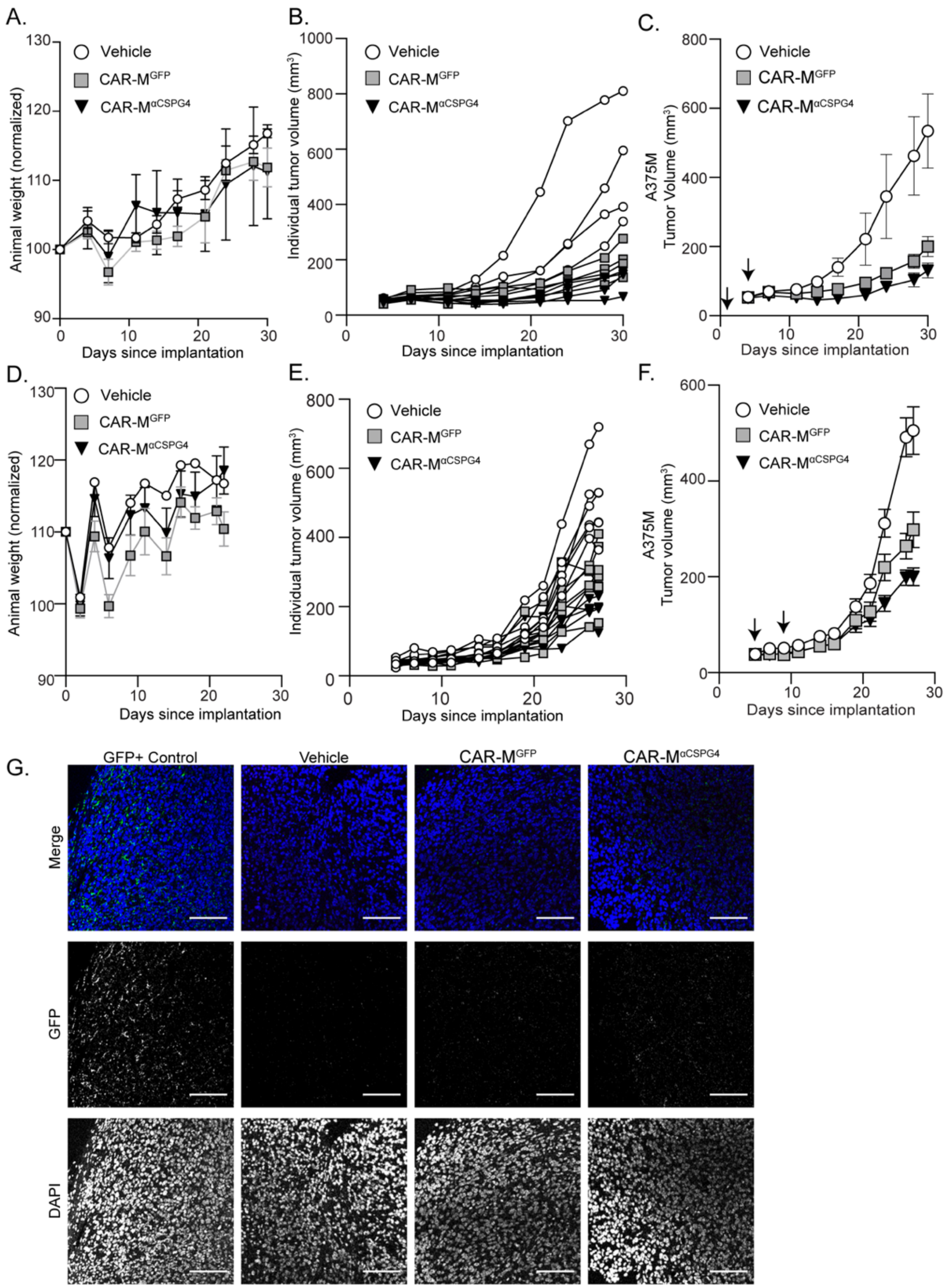


Supplemental Fig 9

**Supplemental Figure 9|** CSPG4-targeting CAR-Ms do not affect mouse weight and do not persist in tumors 5 days after last peritumoral injection. **A-C)** Animal weights (A), individual mouse tumor volumes (B), grouped mouse tumor volumes including vehicle (C) from Figure 6D. **D-F)** Animal weights (D), individual mouse tumor volumes (E), and grouped mouse tumor volumes including vehicle (F) from Figure 6G. **G)** Representative 40X images of immunofluorescent staining for GFP and DAPI of tumors from Figure 6G. As a positive control for GFP staining, a tumor isolated from a GFP+ mouse was used. C, F) Mean +/- SEM. Arrows indicate CAR-M injections.

### Supplemental Video Legends

**Supplemental Video 1|** CSPG4-targeting CAR-Ms fully engulf melanoma cells in 3D. Video of sequential z-planes from confocal images of Fig. 1G – CAR-M$^{\alpha CSPG4(763.74)}$ (green) with internalized A375 cells tagged with Lck-mScarlet (magenta).

**Supplemental Video 2|** CSPG4-targeting CAR-Ms trogocytose melanoma cells in 2D. Timelapse video (maximum intensity projection) of CAR-M$^{\alpha CSPG4(763.74)}$ (green) interacting and trogocytosing/nibbling A375 cells tagged with Lck-mScarlet (magenta). Yellow arrows highlight CAR-M$^{\alpha CSPG4(763.74)}$ actively trogocytosing A375-Lck-mScarlet cell. Images taken every 10 minutes for 18 hours; 7 fps.

**Supplemental Video 3|** CAR-M $^{\alpha CSPG4}$ infiltrate A375-Lck-mScarlet spheroids. Video of a Z-stack showing GFP+ CAR-M$^{\alpha CSPG4}$ (green) inside an A375-Lck-mScarlet spheroid (magenta).

**Supplemental Video 4|** CSPG4-targeting CAR-Ms exhibit increased infiltration of melanoma spheroids compared to control CAR-Ms. Timelapse video of CAR-M$^{GFP}$ (green) that remain on the exterior of A375-H2B-mCherry spheroids (left side) compared to CAR-M$^{\alpha CSPG4}$ which infiltrate A375-H2B-mCherry spheroids (right side). Images taken every 8 hours for 10 days; 7 fps.
